# PD-L1 Restrains the PD-1^hi^Nrp1^lo^TGFβ^+^ Treg to block IL6^+^ Neutrophil tumor infiltration to Suppress Inflammation-driven Colorectal Cancer

**DOI:** 10.1101/2023.10.12.562132

**Authors:** Dakota B. Poschel, John D. Klement, Alyssa D. Merting, Chunwan Lu, Yang Zhao, Dafeng Yang, Wei Xiao, Huabin Zhu, Ponnala Rajeshwari, Michael Toscano, Kimya Jones, Kiran Madwani, Amanda Barrett, Roni J. Bollag, Padraic G. Fallon, Huidong Shi, Kebin Liu

**Author notes:** Correspondence: Kebin Liu, or Huidong Shi,.

## Abstract

PD-L1 functions as a suppressor of T cell activation and colonic inflammation. The consequence of and mechanism underlying these opposite functions of PD-L1 in colorectal cancer remains unknown. We report that global *Cd274* deletion promotes inflammation-driven colorectal tumorigenesis. 16S rRNA and shotgun metagenomic sequencing revealed that loss of host PD-L1 leads to expansion of gut *Ligilactobacillus murinus* and activation of the AhR pathway in tumor-bearing mice. scRNA-seq analysis revealed that PD-L1 regulates PD-1^89^Nrp1^1215^ Treg, IL6^+^ neutrophils, and B cells in colorectal tumor. Treg expresses high level of TGFβ to recruit IL6^+^ neutrophils. IL6 inhibits activation of B and T cells. IL6 blockade or B cell activation via CD40 agonist increases CTL activation and suppresses colon tumor growth in vivo. Our findings determine that PD-L1 functions as a tumor suppressor in the context of inflammation-driven colorectal cancer and the PD-L1/*L. murius*/PD-1^89^Nrp1^1215^TGFβ^+^ Treg/IL6^+^ neutrophils pathway controls host cancer immunosurveillance and colorectal tumorigenesis.

**Key points:** Global deletion of *Cd274* promotes inflammation-driven colorectal tumorigenesis.

Loss of PD-L1 increases gut *L. murinus* and activates AhR pathway in colorectal tumor.

PD-1^hi^TGFβ^+^Nrp1^lo^ Treg recruits IL6^+^ neutrophils to inhibit B and T cell function in colorectal tumor.

*L. murinus* and tryptophan metabolites connect PD-L1 and colorectal tumor immunosurveillance.

**Graphic summary:** 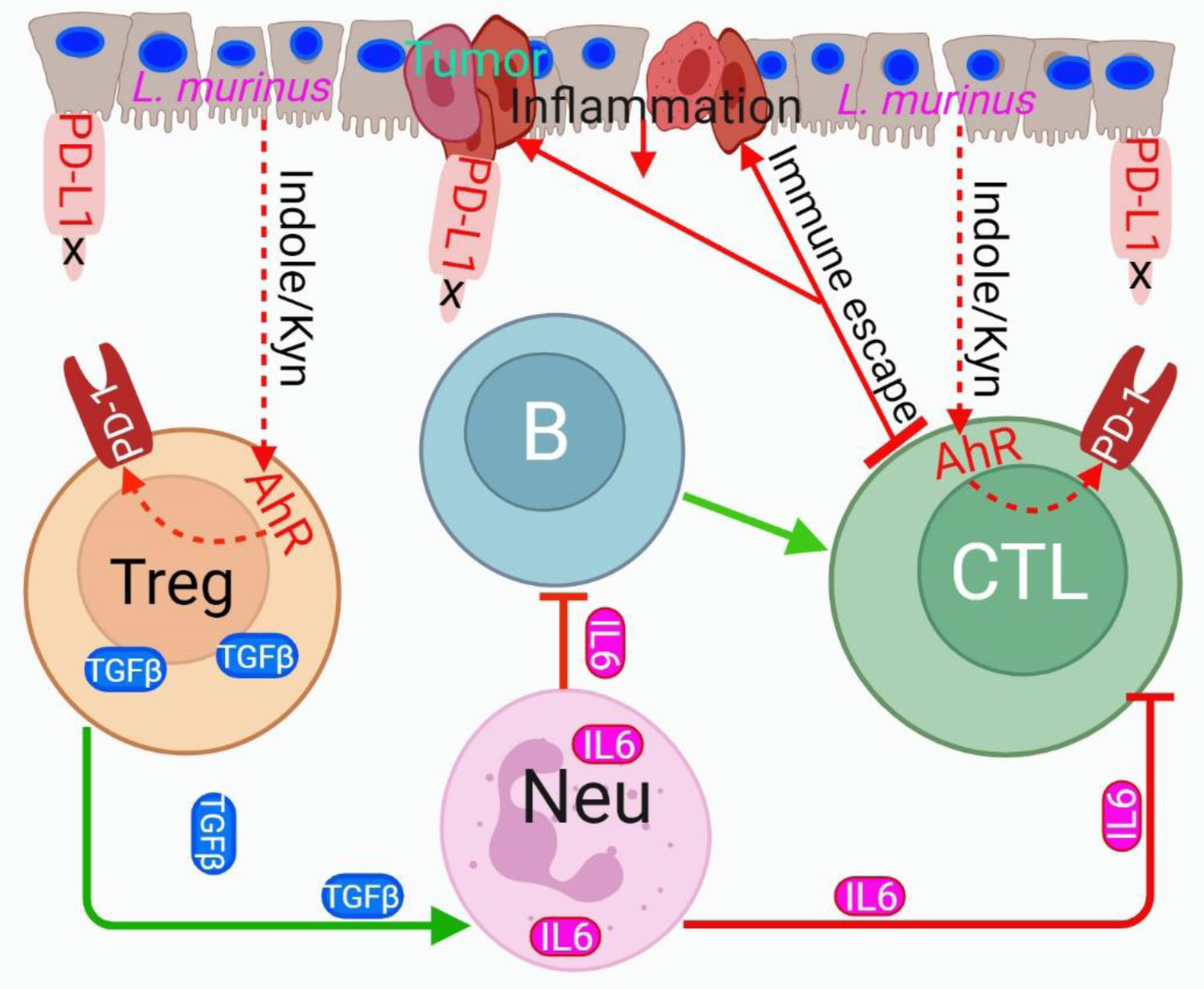

## Introduction

The programmed death-ligand 1 (PD-L1) is a cell surface protein that binds to the programmed cell death protein 1 (PD-1) receptor to inhibit T and myeloid cell activation^1–6^. In human cancer patients and tumor-bearing mice, PD-L1 expression is elevated and engages PD-1 expressed on T cells to activate intrinsic SHP2 to dephosphorylate T cell receptor (TCR) and co-stimulatory receptor CD28 to inhibit T cell activation^7,8^. PD-L1 also binds to PD-1 expressed on myeloid cells to reprogram myeloid cell metabolism to promote differentiation of myeloid-derived suppressor cells^4^ and to repress IRF8-dependent myeloid cell differentiation, resulting in suppression of T cell activation^3^. Furthermore, PD-L1 engages myeloid cell PD-1 to activate the SHP2-type I interferon (IFN)-STAT1-CXCL9 pathway to impair CTL tumor recruitment^5^. The PD-(L)1 pathway therefore acts as an immune checkpoint that negatively regulates T cell activation and tumor recruitment to promote tumor immune escape. PD-(L)1 immune checkpoint inhibitor (ICI) immunotherapy has been developed to re-activate the exhausted and dysfunctional T cells and has shown durable efficacy in human cancer^9–11^. However, human colorectal cancer, except for the small subset of microsatellite instability high (MSI-H) colorectal cancer^9,11^, does not respond to PD-(L)1 ICI immunotherapy^10^.

A unique feature of colorectal cancer is the anatomical location of the tumor to the bacterial neighbors. Billions of commensal bacteria reside in the gut and form the gut microbiota. Dysregulation of gut microbiota causes colonic inflammation and colitis^12,13^, a process that promotes colorectal tumorigenesis^14^. Therefore, it has been hypothesized that the colorectal epithelium is in a continual state of low-grade inflammation due to constant response to the gut microbiome^13,15^. It is thus likely that certain colorectal cancer could be the consequence of microbiome-induced inflammation^12,13,16^. However, what controls this gut microbiota-mediated continuous chronic colonic inflammation is unknown.

Approximately 85-90% human colorectal cancer is the microsatellite stable (MSS) type, and only about 10-15% human colorectal cancer is the MSI-H subtype^17,18^. Paradoxically, MSS human colorectal cancer expresses weak to no PD-L1, whereas MSI-H human colorectal cancer expresses high level of PD-L1^19,20^, thereby suggesting that PD-L1 might functions overall as a tumor suppressor in the context of MSS human colorectal cancer. This notion is consistent with the clinical finding that colonic inflammation and colitis is an immune-related adverse effect of the PD-(L)1 ICI immunotherapy^21–23^. Furthermore, administrating Fc-fused PD-L1 protein abrogates colonic inflammation and colitis in experimental colitis mouse models^24^. The most definitive proof of PD-L1 function in suppression of colonic inflammation came from studies with PD-L1 KO mice showing that inflammation-inducing agent causes significantly more severe colonic inflammation and colitis in PD-L1 KO mice than in WT mice^25^. PD-L1 therefore not only functions as an immune checkpoint to promote tumor immune evasion, but also as a suppressor of colonic inflammation that might prevent colorectal tumorigenesis. However, the consequences of and mechanisms underlying these two opposite functions of PD-L1 in colorectal cancer development are incompletely understood.

Given that colon is in a continual state of low-grade inflammation^15^ and PD-L1 plays an essential role in suppression of colonic inflammation^24,25^. It is therefore possible that loss of PD-L1 in colorectal cancer shifts the balance of PD-L1 functions between tumor immune evasion promotion and colonic inflammation suppression towards enhancing colorectal tumorigenesis. PD-L1 therefore functions as a tumor suppressor in the context of colorectal cancer. We aimed at testing this hypothesis. Our findings determine that loss of host PD-L1 increase gut *Lactobacillus* and PD-1^hi^ Nrp1^lo^Treg cells that produces TGFβ to recruit IL6^+^ neutrophils to suppress T and B cell activation to promote sporadic and inflammation-driven colorectal tumorigenesis. The PD-L1/*Lactobacillus*/PD-1^hi^Nrp1^lo^TGFβ^+^Treg/IL6^+^ neutrophils pathway controls B and T cell-dependent host cancer immunosurveillance and colorectal tumorigenesis.

## Results

### Global PD-L1 deficiency increases sporadic and inflammation-driven colon tumorigenesis

PD-L1 is expression in both tumor cells and immune cells ^1,2,26^. To determine the cell types that express PD-L1 in human colorectal cancer, human colorectal tumor tissues were prepared into single cells and stained with CD45- and PD-L1-specific antibodies by immunohistochemical analysis. Consistent with what was reported in human colorectal cancer patients ^19,20^, PD-L1 is expressed in both CD45^+^ immune cells and tumor cells (Fig. S1A). We then extract scRNA-seq dataset of human colon cancer patients^27^in the single cell portal (Brad Institute) and analyzed PD-L1 expression profile in human colon cancer in the single cell level. Low level of PD-L1 is detected in subsets of immune cells, including B, T, NK, and ILCs, and stromal cells. Tumor cells are the most heterogeneous cell population that express PD-L1. Myeloid cells expressed the highest level of PD-L1 (Fig. S1B).

To determine the relative functions of cell type-specific PD-L1, we made use of the azoxymethane (AOM) and dextran sodium sulfate (DSS)-induced sporadic and inflammation-driven colorectal cancer mouse model^28^. Analysis of colorectal tissues of tumor-free mice determined that PD-L1 protein is located on the surfaces of colorectal epithelial cells and colorectal resident immune cells (Fig. 1A). In the mouse colorectal tumor tissue, PD-L1 protein is present on the surfaces of both tumor cells and colorectal tumor-infiltrating immune cells (Fig. 1B). Further analysis revealed that PD-L1 protein is also present on myeloid cell surface (Fig. 1B). Taken together, these findings determine that this sporadic and inflammation-driven colorectal tumor mouse model resembles human colorectal cancer in PD-L1 expression profiles.

**Figure 1.**
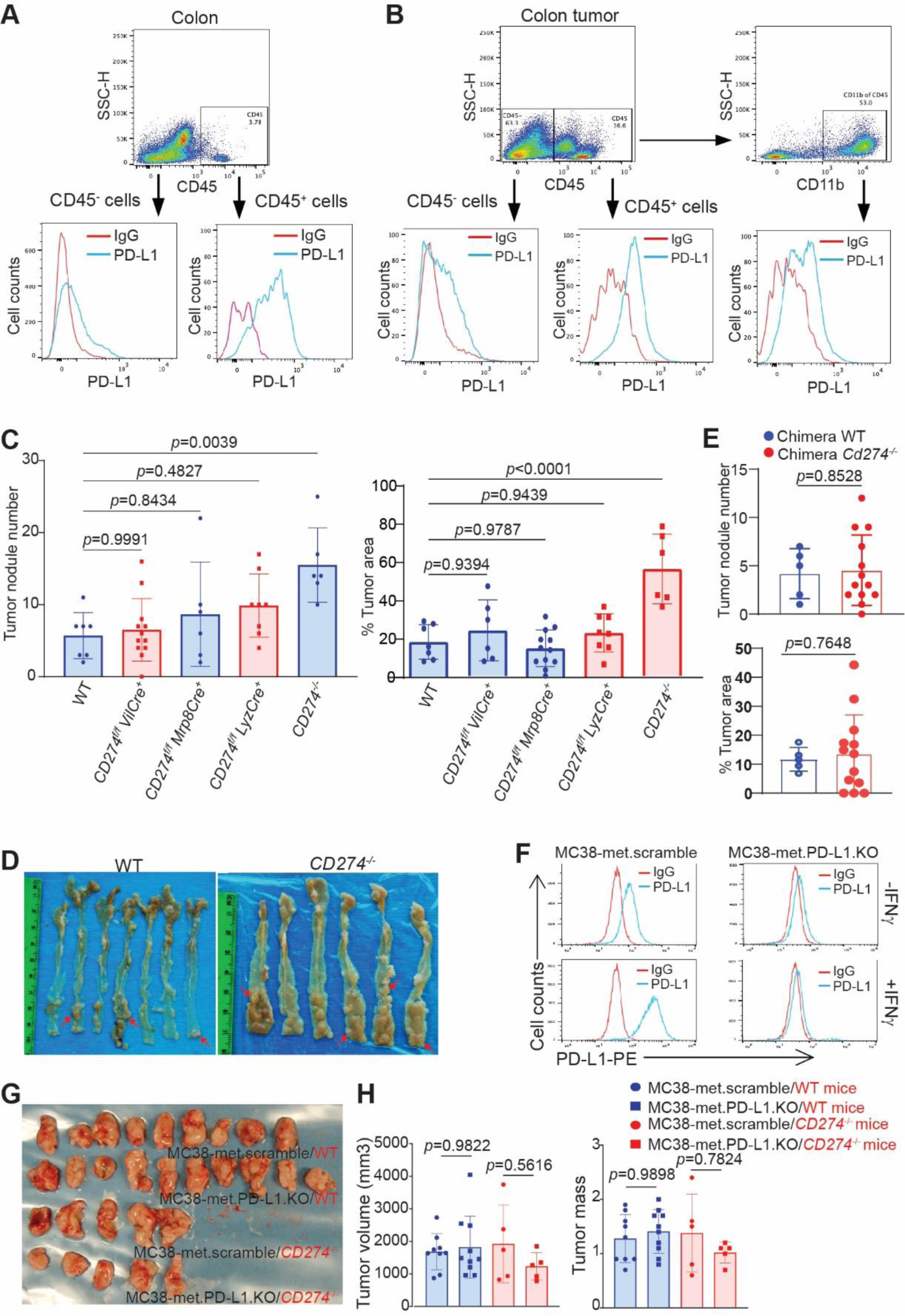
PD-L1 functions as a tumor suppressor in inflammation-driven colorectal tumor-bearing mice. **A**. PD-L1 expression in colon epithelial cells (CD45^-^) and colon resident immune cells (CD45^+^) of tumor-free mice. Representative data of one of three mice. **B**. PD-L1 expression in tumor cells (CD45^-^), tumor-infiltrating immune cells (CD45^+^), and myeloid cells (CD11b^+^) in tumors from colon tumor-bearing mice. Representative data of one of three tumor-bearing mice. **C**. Colorectal tumor nodule number and tumor size in WT mice and mice with *Cd274* deletion in the indicated cell types. **D**. Colorectal tumorigenesis in WT mice and mice with *Cd274* global deletion. Shown are tumor-bearing colon. **E**. Colorectal tumor nodule number and tumor size in PD-L1 KO chimera mice and WT chimera control mice. **F**. Phenotypes of WT (MC38-met.scramble) and PD-L1 KO (MC38-met.PD-L1.KO) tumor cells lines. **G & H**. MC38-met.scramble and MC38.met.PD-L1.KO cells were injected subcutaneously to WT and PD-L1 KO mice. Shown are tumor image (left panel) and quantification of tumor size and weight (right panel).

Myeloid cell PD-L1 has been shown to promote tumor immune evasion in transplanted cutaneous tumor mouse models^29–33^. To determine myeloid PD-L1 function in spontaneous colorectal tumorigenesis under pathophysiological conditions, we created mice with PD-L1 deletion only in myeloid cells by crossing *Cd274* floxed mouse^34^ to *Lyz*-*Cre* and *Mrp8*-*Cre* mice, respectively. *Lyz-Cre* generated mice with PD-L1 deletion only in myeloid cells and *Mrp8-Cre* mice generate mice with PD-L1 deletion only in neutrophils. Colorectal tumors were then induced in these mice. Analysis of the colorectal tissues determined that knocking out PD-L1 in myeloid cells or neutrophils has no significant effect on colorectal tumorigenesis (Fig. 1C). We then sought to knock out PD-L1 in all immune cells using adoptive transfer of bone marrow (BM) cells from the PD-L1 KO mice^35^ to lethally irradiated WT recipient mice to create PD-L1 KO chimera mice^36^. Similar to what was observed in myeloid and neutrophil PD-L1 KO mice, deletion of PD-L1 in immune cells has no significant effect on colorectal tumorigenesis (Fig. 1E). It has also been reported that tumor cell PD-L1 alone is sufficient to promote tumor immune evasion in a transplanted colon tumor mouse model^37^. We then created mice with PD-L1 deletion only in colorectal epithelial cells. PD-L1 deletion in colorectal epithelial cells also has no significant effect on colorectal tumorigenesis (Fig. 1C). It has also been reported that both tumor cells and host PD-L1 are required for suppression of host anti-tumor immune response^35^, our above findings suggest that colon epithelial cell/tumor cell PD-L1 and host immune cell PD-L1 may compensate each other to promote tumor immune evasion. To test this hypothesis, we induced colorectal tumor in mice with global PD-L1 deletion. Strikingly, instead of the expected decreased tumorigenesis, global deletion of PD-L1 significantly increased colorectal tumor nodule number and tumor size as compared to WT mice in this sporadic and inflammation-driven colorectal tumorigenesis mouse model (Fig. 1C & D).

To determine whether the above finding is colonic inflammation-driven colon tumor-specific, we treated WT and PD-L1 KO mice with AOM only for five times weekly and analyzed colon days later. The colon tissues were analyzed by two board-certified pathologists. No tumor formation was observed in the colon of the WT and PD-L1 KO mice (Fig. S2A). To further strengthen this finding, we then used PD-L1 KO mouse colon tumor cell lines (Fig. 1F) and transplanted WT and PD-L1 KO tumor cells to WT and PD-L1 KO mice, respectively. The subcutaneous tumor is relatively distant from the colon and thus minimizes the direct effect of colonical inflammation. In this subcutaneous colon tumor mouse model, we observed no effect of PD-L1 on colon tumor growth in both WT and PD-L1 KO mice (Fig. 1G & H), indicating that PD-L1 function in promoting colorectal tumorigenesis is colon organ-specific. Taken together, our findings indicate that PD-L1 functions as a suppressor of inflammation-driven colorectal tumorigenesis in mice.

### Host PD-L1 controls gut microbiome and suppresses colonic inflammation

PD-L1 knock out mice exhibit severe colonic inflammation and inflammation-associated colitis when treated with the inflammatory agent^25^. PD-L1 protein inhibits colonic inflammation and colitis in mice^24^. A major adverse effect of PD-(L)1 ICI immunotherapy in human cancer patients is ulcerative colitis ^22,38–41^. PD-L1 therefore plays an essential role in protecting colorectal epithelium from colonic inflammation. Consistent with these findings, we observed that PD-L1 KO mice exhibit significant greater body weight loss (Fig. S2B) and shortened colon (Fig. S2D) as compared to WT mice when treated with AOM and DSS. However, there is no significant difference in mouse survival after AOM-DSS treatment between WT and PD-L1 KO mice (Fig. S2D). Histological analysis of colorectal tissues revealed that PD-L1 KO mice exhibit greater degree of colonic inflammation as measured by inflammation score (Fig. 2E)^42^. Colorectal adenoma and carcinoma developed in both WT and PD-L1 KO mice at day 21 after AOM-DSS treatment with a higher grade of tumor burden in the PD-L1 KO mice (Fig. 2E). At day 79, colonic inflammation decreased to a lower level in both WT and PD-L1 KO, and both WT and PD-L1 KO mice have high-grade adenoma and carcinoma (Fig. 2E). However, PD-L1 KO mice have significantly more tumor nodules and bigger tumor nodule size than WT mice (Figs.1C-D). These observations indicate that loss of host PD-L1 results in increased colonic inflammation to promote inflammation-driven colorectal tumorigenesis in mice.

**Figure 2.**
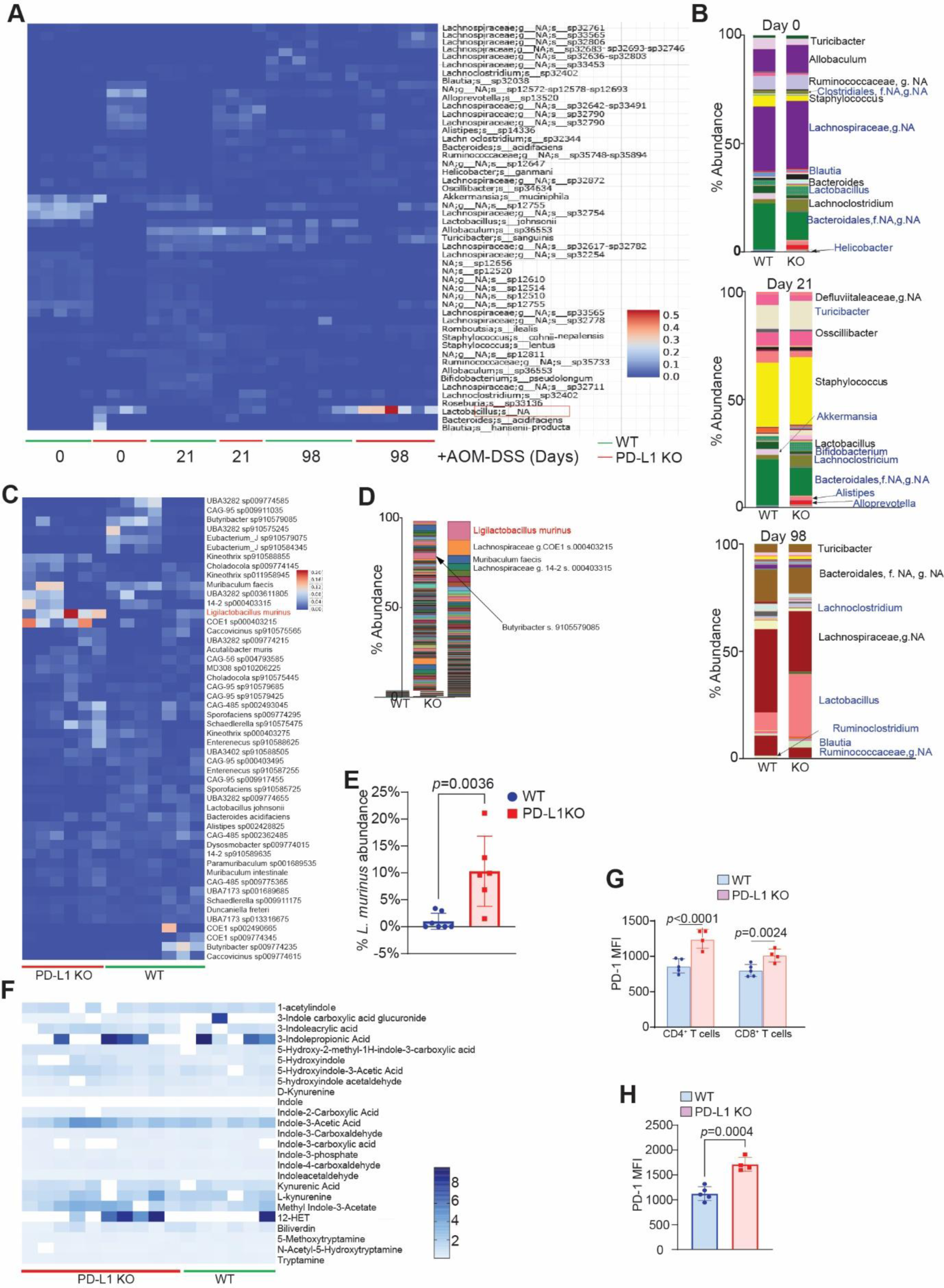
Host PD-L1 suppresses *Ligilactobacillus murinus* in the gut in colon tumor-bearing mice. **A.** Heatmap showing abundance and patterns of gut microbiota genus in WT and PD-L1 KO mice as indicated. **B**. Percentage of gut microbiota in WT and PD-L1 KO mice at the indicated stage of colon tumor development as shown in A. **C**. Heatmap showing gut microbiome species in WT and PD-L1 KO mice. **D**. Percentage of gut microbiome species in WT and PD-L1 KO mice as shown in C. **E**. *L. murinus* abundance. **F**. Heatmap showing serum tryptophan metabolite level in tumor-bearing WT and PD-L1 KO mice. **G**. PD-1 level in the T and myeloid cells in tumor-bearing WT and PD-L1 KO mice. **H**. PD-1 expression level in Treg cells in in tumor-bearing WT and PD-L1 KO mice.

Emerging experimental data indicate that the gut microbiota plays a critical role in colonic inflammation and host anti-tumor immune response^12,16,43,44^. To determine whether host PD-L1 regulates gut microbiome, we collected fecal specimens from WT and PD-L1 KO mice at different time points after AOM-DSS treatment and performed16S rRNA gene sequencing. Distinct microbe profiles were observed in WT and PD-L1 KO mice and AOM-DSS altered the microbiota differently in WT and PD-L1 KO mice over the course of colorectal tumorigenesis (Fig. 2A & B). Analysis of the differential microbiome of known genus revealed that knocking out PD-L1 in the host decrease abundance of genus *Bacteroides* in the gut and this differential *Bacteroides* pattern persists in the gut in the early stage after AOM-DSS treatment. However, *Bacteroides* population diminished at the late stage after abundant colorectal tumor formed in both WT and PD-L1 KO mice (Fig. 2B). The PD-L1 KO mice have a significantly higher level of genus *Lactobacillus* than WT mice and *Lactobacillus* remained higher in the PD-L1 KO mice than WT mice in the early stage of AOM-DSS treatment. Strikingly, gut *Lactobacillus* abundance increased dramatically in the colorectal tumor-bearing mice and its abundance is significantly higher in PD-L1 KO mice than in WT mice. *Lactobacillus* genus accounts for approximately 29.13% and 8.58% of the gut microbes in the colorectal tumor-bearing PD-L1 KO and WT mice, respectively (Fig. 2B).

To further define the species in *Lactobacillus* genus, we then performed a shotgun metagenomic sequencing of the fecal samples. Consistent with the 16S rRNA gene sequencing data, metagenomic sequencing analysis determined that *Ligilactobacillus murinus* (formerly *Lactobacillus murinus*) is the most dominant species in the *Lactobacillus* genus as identified by the 16S rRNA sequencing and accounts for 10.31% of the total gut microbiomes in tumor-bearing PD-L1 KO mice. In contrast, *L. murinus* only accounts for 1.04% of the total gut micromes in tumor-bearing WT mice (Figs. 2C-E). Our finding determine that host PD-L1 functions to suppress gut *L. murinus* in colorectal tumor-bearing mice.

*L. murinus* is known to metabolize dietary tryptophan to indoles and kynurenine to activate aryl hydrocarbon receptor (AhR) in tumor-associated macrophage to suppress CTL anti-tumor immunity^44–48^. We next performed metabolomics analysis of mouse serum. Compared to WT mice, the most significant change of metabolites in PD-L1 KO colorectal tumor-bearing mice is amino acid and its metabolites (Fig. S3A). Targeted analysis of the tryptophan metabolites level varies between individual mice. However, a higher percentage of PD-L1 KO mice have higher serum level tryptophan metabolites including 3-indopropionic acid, indole-3-acetic acid, indole-3-phosphate, L-kynurenine, and 12-HETE, as compared to WT tumor-bearing mice (Fig. 2F).

The activated AhR is a cytosolic transcription factor that translocate to the nucleus to activate *Pdcd1* transcription to increase PD-1 protein level on T cells^49^. PD-1 in T cells is therefore a response signature of the activated AhR pathway^49^. We then analyzed PD-1 level in T cells in tumor-bearing mice. Flow cytometry analysis indicates that, as expected, PD-1 level is higher in CD4^+^ and CD8^+^ T cells in the tumor-bearing PD-L1 KO as compared to WT mice (Fig. S3B, Fig. 2G). Furthermore, PD-1 protein level is also significantly higher on Treg cells in tumor-bearing PD-L1 KO mice than in WT mice (Fig. 2H). The percentages of PD-1^+^ T cells are also significantly different in spleens between WT and PD-L1 KO mice (Fig. S3B & C). These observations indicate that loss of host PD-L1 leads to an increase in abundance of gut *L. murinus,* its tryptophan metabolites, and activation of the AhR pathway in colorectal tumor-bearing mice.

### A high-resolution cellular landscape of PD-L1-deficient mouse colorectal tumor

Our above findings determine that, despite its well-documented function in promotion of tumor immune evasion, PD-L1 functions as a tumor suppressor in the context of inflammation-driven colorectal tumor mouse model. To elucidate the molecular and cellular mechanisms underlying PD-L1 function in colonic inflammation and colorectal tumorigenesis, we analyzed the colorectal tumor nodules dissected from tumor-bearing WT and PD-L1 KO mice by single-cell RNA sequencing (scRNA-seq)(Fig. 3A). A total of 24,000 single cells passed quality control and were annotated using canonical lineage markers. This high-level annotation was further confirmed using published gene signatures^50^. Uniform manifold approximation and projection (UMAP) visualization identified 26 unique cell clusters/population in the colorectal tumor tissues (Fig. 3B) with unique gene signatures (Fig. S4A). PD-1 transcripts are presented in two cell clusters (Fig. 4B). Consistent with the increased PD-1 protein level in T cells of PD-L1 KO mice than in WT mice (Figs. 2G & H), PD-1 transcript level is also higher in in these two clusters in PD-L1 KO mice than in WT mice (Fig. S4B). Only one exon in the *Cd274* gene was deleted in the PD-L1 KO mice^35^ and PD-L1 transcripts are detected in 18 cell clusters. PD-L2 transcripts are detected in the same 18 cell clusters as PD-L1 (Fig. S4B). Tumors from the WT and PD-L1 KO mice exhibit similar cellular clusters (Fig. 3B). Nearest neighbor clustering identified 12 cell populations (Fig. 3C) and each cell population has unique gene signatures (Fig. 3D). Among these 12 cell populations, B cell abundance decreased dramatically whereas neutrophils and T cells increased. Further analysis of the immune cell population validated the decreased B cells and increased neutrophils and T cell populations in colorectal tumors from PD-L1 KO mice (Fig. 3E). PD-1 transcript is primarily detected in NK and T cells, PD-L1 and PD-L2 are expressed in all major immune cell subpopulations (Fig. S5A & B). PD-1 level, the AhR response signature^49^, is higher in NK/T cells in PD-L1 KO tumor-bearing mice than WT tumor-bearing mice. To further determine the differential AhR activation, we analyze AhR expression levels in the cell subpopulation. AhR expression level is higher in monocytes, macrophages, and DCs (Fig. S5C). We then used the 166-gene AhR pathway activation signature^51^ and determined that AhR pathway activation level is greater in stromal cells, monocytes, macrophages, DCs, and mast cells in colorectal tumor in PD-L1 KO mice than in WT mice (Fig. S5D). Our finding thus determine that decreased B cells, increased neutrophils and T cells, and AhR pathway activation are hallmarks of inflammation-driven colon cancer in mice.

**Figure 3.**
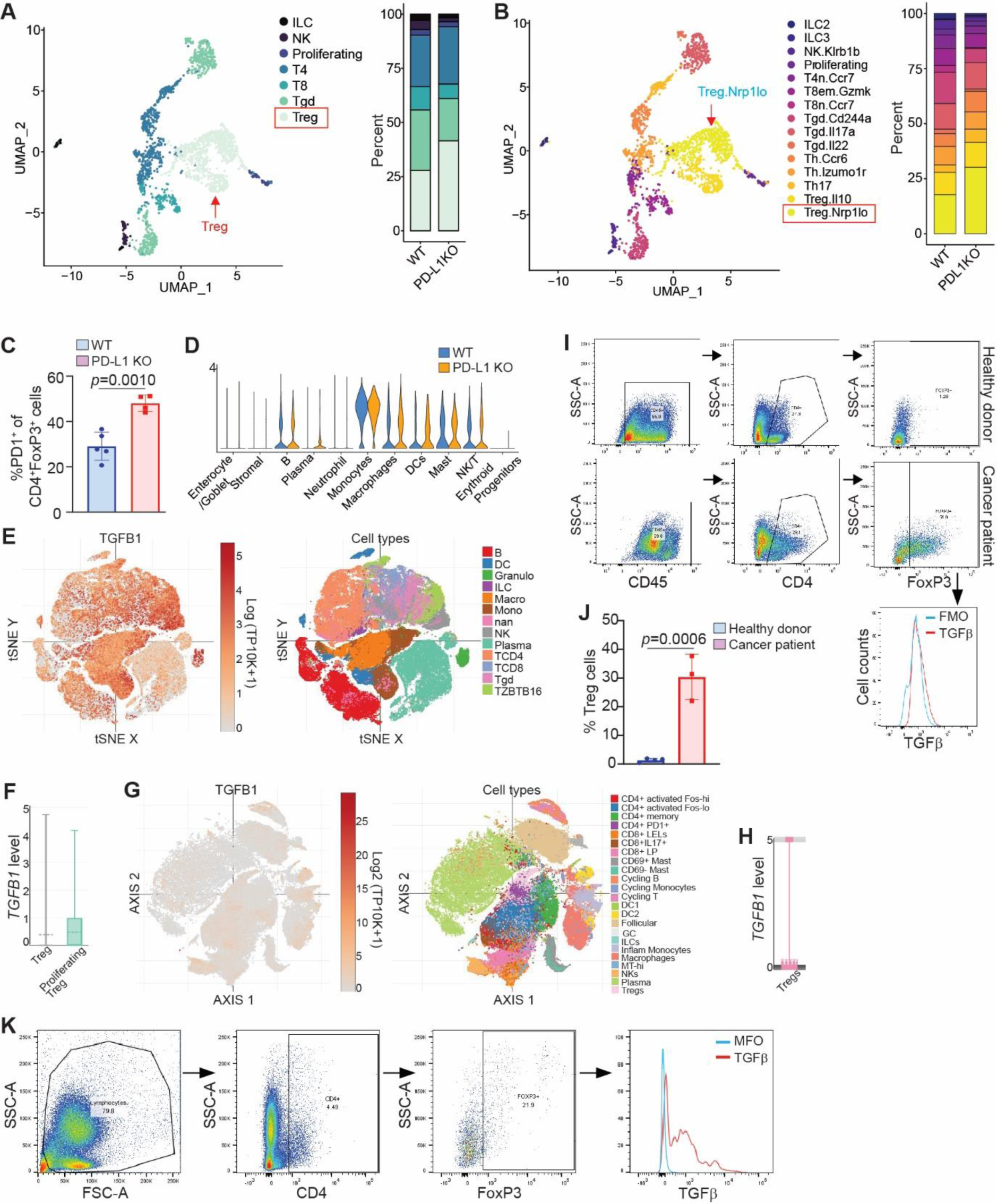
Tumor cell and immune cell landscape of WT and PD-L1 KO mouse colon tumor. **A**. scRNA-seq experimental scheme. **B**. UMAP plot of all cells isolated from colon tumor, colored by identified cell clusters (right panel). The cell cluster overlap of tumors by tumor-bearing mouse genotypes are shown at the right panel. **C**. UMAP plot (left) and barplot (right) of identities of the cell subpopulation of total colon tumor cells as shown in B. **D**. Heatmap of top 5 transcripts as indicated for each cell subpopulation as shown in C. **E**. UMAP plot (left) and barlpot (right) of identities of the subpopulation of CD45+ cells in the colon tumor.

**Figure 4.**
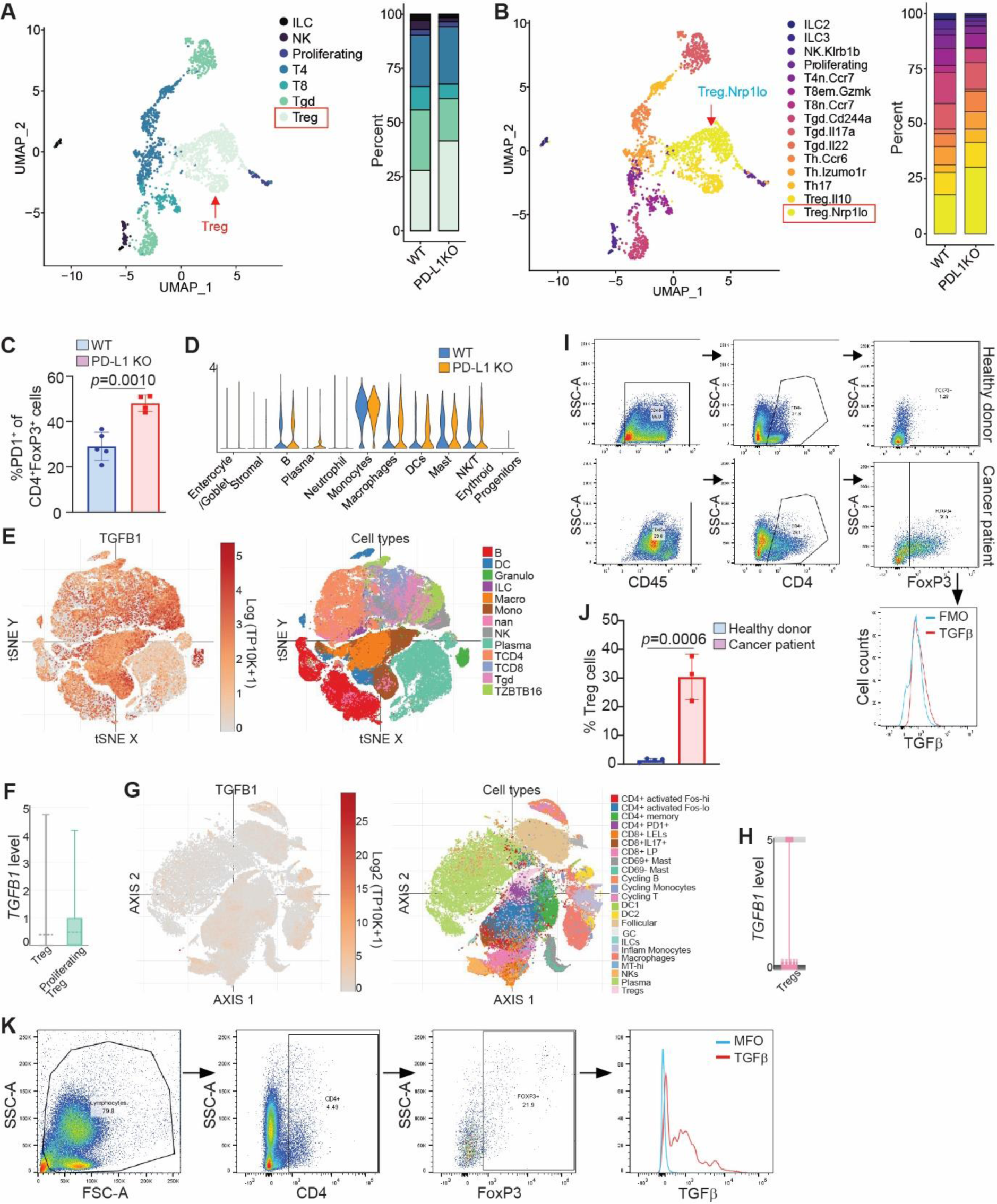
PD-L1 suppresses TGFβ-expressing Nrp1^lo^ Treg cells accumulation in colon tumor. **A**. UMAP projection (left) and barplot (right) of T, NK, and ILC in colon tumor. Treg, T regulatory cells, Tgd, γδ T cells, T8, CD8^+^ T cells; T4, CD4^+^ T cells; Proliferating, Proliferating cells; NK, NK cells; and ILC, innate lymphoid cells. **B**. UMAP projection (left) and barplot (right) of subpopulations of T, NK, and ILCs. **C**. Quantification of PD-1^+^ Treg cells in WT (n=5) and PD-L1 KO (n=4) tumor-bearing mice. **D**. Violin plot of *Tgfb1* transcripts in the indicated cell populations in colon tumor from WT and PD-L1 KO mice. **E**. UMAP plots of *TGFB1* transcripts (left panel) and immune cells (right) of human colon tumor. Human colorectal cancer dataset were retrieved from the single cell portal (GEO:GSE178341). **F**. Box plot of *TGFB1* transcript level in the indicated cell subpopulation. **G**. UMAP projection of *TGFB1* (left) and immune cell subpopulation (right) of human colon tumor as shown in E. **H**. Box plot of *TGFB1* transcript level in Treg cells. **I**. TGFβ expression level in Treg cells in the peripheral blood of healthy donors (n=3) and colorectal cancer patients (n=3). Shown is gating strategy and TGFβ MFI. **J**. Quantification of Treg cell levels in the peripheral blood of healthy donors and colorectal cancer patients. **K**. Human colon tumor tissue was analyzed for Treg. Shown are gating strategy and TGFβ level in Treg cells.

### PD-L1 suppresses Nrp1^lo^ Treg cell accumulation and TGFβ production

Given the function of PD-L1 as a potent inhibitor of T cells in the tumor microenvironment^1–3,5^, our above observation that PD-L1 KO tumor has increased T cell accumulation is expected. We then analyzed T cell subpopulations in the single cell level. Clustering analysis demarcated populations of CD4^+^, CD8^+^, γδ T, and Treg cells. Surprisingly, the levels of CD4^+^ and CD8^+^ T cells are not significantly increased in the PD-L1 KO colorectal tumor (Fig. 4A&B, Fig. S6). Furthermore, Treg and γδ T cells are the major populations of T cells and Treg cell level increased dramatically in the PD-L1 KO colorectal tumor (Fig. 4A&B). Among the Treg cells, the majority is Nrp1^lo^ subset (Fig. 4B). As reported in the literature, gut *Lactobacillus* produces indoles and kynurenine which are known to activate AhR^44,45^. Activated AhR is known to induce Treg cell accumulation^52–55^. We observed that AhR pathway is activated in stromal cells, monocytes, macrophages, DCs, and mast cells in colorectal tumor in PD-L1 KO mice than in WT mice (Fig. S5D). Consistent with this phenomenon, PD-1 level is significantly higher in Treg cells in tumor-bearing PD-L1 KO mice than in WT mice (Fig. 4C). Our findings thus indicate that host PD-L1 suppresses the PD-1^hi^Nrp1^lo^Treg cells in inflammation-driven colorectal tumor.

TGFβ is a cytokine that not only regulates Treg differentiation but is also expressed in Treg cells^56,57^. Analysis of the immune cell subsets revealed that TGFβ transcript is expressed in various cell types in WT and PD-L1 KO mouse colon tumors (Fig. 4D). Analysis of human colorectal cancer patient scRNA-seq datasets^27^ validated that *TGFB1* transcript is expressed in various immune cell populations (Figs. 4E & G). Further analysis revealed that *TGFB1* is highly expressed in Treg cells in human colorectal cancer patients (Fig. 4F & H). Analysis of peripheral blood samples determined that TGFβ is expressed in Treg cells in colorectal cancer patients (Fig. 4I) and Treg level is significantly higher in colorectal cancer patients than in healthy donors (Fig. 4J). Analysis of human colorectal tumor indicates that TGFβ is expressed in tumor-infiltrating Treg cells (Fig. 4K). Our findings determine that loss of host PD-L1 results in accumulation of PD-1^hi^Nrp1^lo^TGFβ^+^Treg cells.

### PD-L1 deficiency increases neutrophil accumulation and IL6 production

Neutrophils are a highly heterogeneous population in human cancer patients and tumor-bearing mice^58–60^. Our above finding revealed that loss of PD-L1 leads to expansion of neutrophils in colorectal tumor. To determine whether PD-L1 regulates neutrophil subpopulation differentiation, neutrophil subsets were further identified. Five unique neutrophil clusters were identified, and each subset has unique gene signatures (Fig. 5A & B). However, PD-L1 deficiency did not change neutrophil subset clusters (Fig. 5B & C), indicating that PD-L1 deficiency leads to expansion of the total neutrophils in colorectal tumor (Fig. 3C), but not unique subsets of neutrophils. Analysis of spleen cells of the AOM-DSS-induced colorectal tumor validated the significant increase of neutrophils in mice with global *Cd274* deletion (Fig. 5D) and in the chimera mice (Fig. 5E). However, mice with *Cd274* deletion only in colorectal epithelial cells or myeloid cells have similar levels of neutrophils as the WT mice (Fig. 5D).

**Figure 5.**
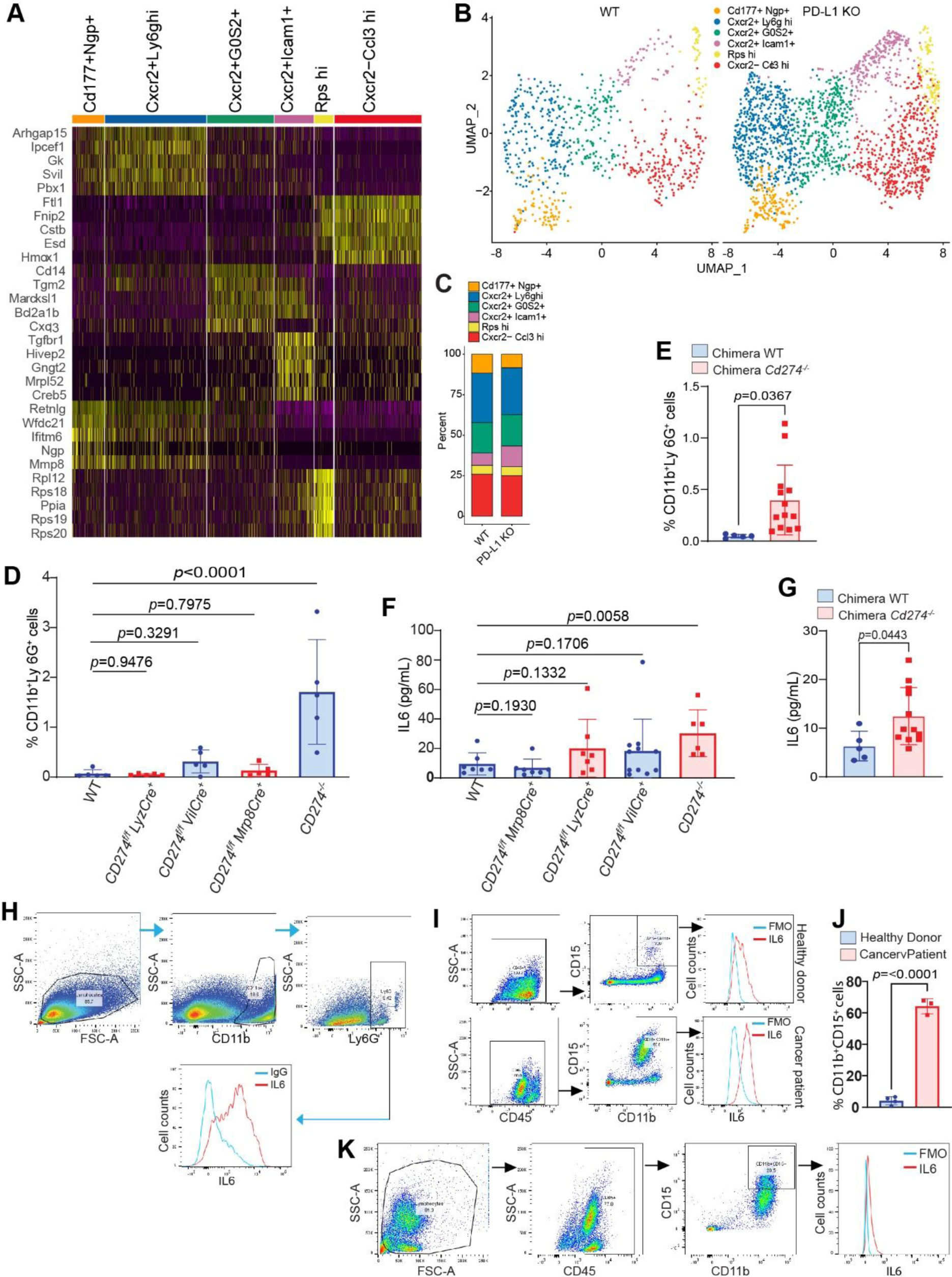
Neutrophils express high level of IL6 in mouse colon tumor. **A**. Dimensional heatmap of top ten most variably expressed genes in each neutrophil subpopulation as indicated. **B & C**. UMAP projection (B) and barplot (C) of neutrophil subpopulations in colon tumor of WT and PD-L1 KO mice. **D**. Quantification of neutrophils in spleens of WT and the indicated tissue-specific and global PD-L1 KO mice by flow cytometry. **E**. Quantification of neutrophils in spleens of WT and PD-L1 KO chimera mice. **F.** IL6 protein level in serum of WT and the indicated tissue-specific and global PD-L1 KO mice. **G**. IL6 protein level in serum of WT and PD-L1 KO chimera mice. **H**. IL6 level in neutrophils of the orthotopic CT26 colon tumor as analyzed by flow cytometry. **I**. IL6 protein level in neutrophils in the peripheral blood of healthy donors (n=3) and colorectal cancer patients (n=3). Shown are gating strategy and IL6 MFI. **J**. Quantification of neutrophil levels in the peripheral blood of healthy donors and colorectal cancer patients as shown in I. **K**. IL6 protein level in human colon tumor. Shown are gating strategy and IL6 MFI in the neutrophils.

Measurement of serum of the tumor-bearing mice shows that IL6 protein level is significantly elevated in the tumor-bearing PD-L1 KO mice and chimera mice as compared to the WT mice (Fig. 5F & G). The level of inflammatory cytokine IL17 is not significantly different between WT and PD-L1 KO mice (Fig. S7). The levels of IFNγ and TNFα, two cytokines produced by activated T cells, are also not significantly different between WT and PD-L1 Ko mice (Fig. S7). Only the levels of T cell chemoattractant Cxcl1, Cxcl9, and Cxcl10 are significantly different between WT and PD-L1 KO tumor-bearing mice (Fig. S7). Consistent with the neutrophil profiles, tumor-bearing mice with *Cd274* deletion only in colorectal epithelial cells or myeloid cells have no significant change in IL6 protein level in the peripheral blood (Fig. 5F). These findings establish a correlation between neutrophil accumulation level and IL6 protein level in the WT and PD-L1-deficient tumor-bearing mice, suggesting that neutrophils are major producers of IL6 in the colorectal tumor-bearing mice. To test this hypothesis, we injected colon tumor cells to mouse cecal wall to establish orthotopic colon tumor. Analysis of the colon tumor identified a distinct population of neutrophils, and the majority of the neutrophils express IL6 (Fig. 5H). Analysis of human peripheral blood samples revealed that CD11b^+^CD15^+^ neutrophils express IL6 (Fig. 5I) and neutrophils are much more abundant in colorectal cancer patient blood in in healthy donor blood (Fig. 5J). Analysis of human colon tumor tissues revealed that tumor-infiltrating neutrophils express high level of IL6 (Fig. 5K). We therefore conclude that PD-L1 deficiency results in expansion of neutrophils in colorectal tumor and tumor-infiltrating neutrophils are major producers of IL6 protein.

### Treg cells recruit IL6^+^ neutrophils via secreting TGFβ

Although TGFβ plays a major role in Treg cell differentiation, emerging experimental data indicate that Treg cells also produce TGFβ^61–63^. Because TGFβ is elevated in the early stage and IL6 protein level is high at the late stage of the tumor development, it is possible that Treg cells may use TGFβ to recruit neutrophils. To test this hypothesis, we performed an in vitro neutrophil recruitment assay. Naïve T cells were purified from mouse spleens and cultured in vitro to induce differentiation into Treg cells. The in vitro differentiated Treg is mostly Nrp1^lo^ and express high level of TGFβ (Fig. 6A). The cultured Treg cells secret TGFβ (Fig. 6B). Neutrophils were then purified from tumor-bearing mice and cultured in transwells in a culture plate in the absence or presence of TGFβ protein. Analysis of neutrophil migration through the transwells shows that TGFβ protein significantly increased neutrophil migration (Fig. 6C). Tumor-associated neutrophils are known to have potent suppressive activity against T cell activation ^64–67^. To determine whether neutrophils suppress T cell activation through IL6, we first co-cultured neutrophils with the antigen-specific 2/20 CTL line and analyzed T cell proliferation. Neutrophils inhibited T cell proliferation in a cell dose-dependent manner (Fig. 6D). To determine whether neutrophils suppress T cell proliferation through IL6, IL6 neutralization mAb was then added to the co-culture. IL6 blockade increased T cell activation in the neutrophil and T cell co-culture (Fig. 6E). To further determine IL6 function in T cell activation, we cultured purified splenic T cells in anti-CD3 and anti-CD28-coated plates in the presence of IL6. Analysis of T cell proliferation indicates that IL6 inhibited T cell activation (Fig. 6F). A hallmark of the PD-L1 deficient colorectal tumor-bearing mice is decreased B cell level (Fig. 3C & E). IL6 is known to inhibit B cells under inflammatory vonditions^68^. To determine whether IL6 also inhibits B cell activation, we purified B cells from mouse spleens and activated B cells with lipopolysaccharides in the absence or presence of IL6 protein. Analysis of B cell proliferation revealed that IL6 significantly inhibited B cell activation (Fig. 6G). Taken together, our findings thus indicate that Treg cells produces TGFβ to recruit neutrophils and neutrophils secret IL6 to inhibit activation of B and T cells.

**Figure 6.**
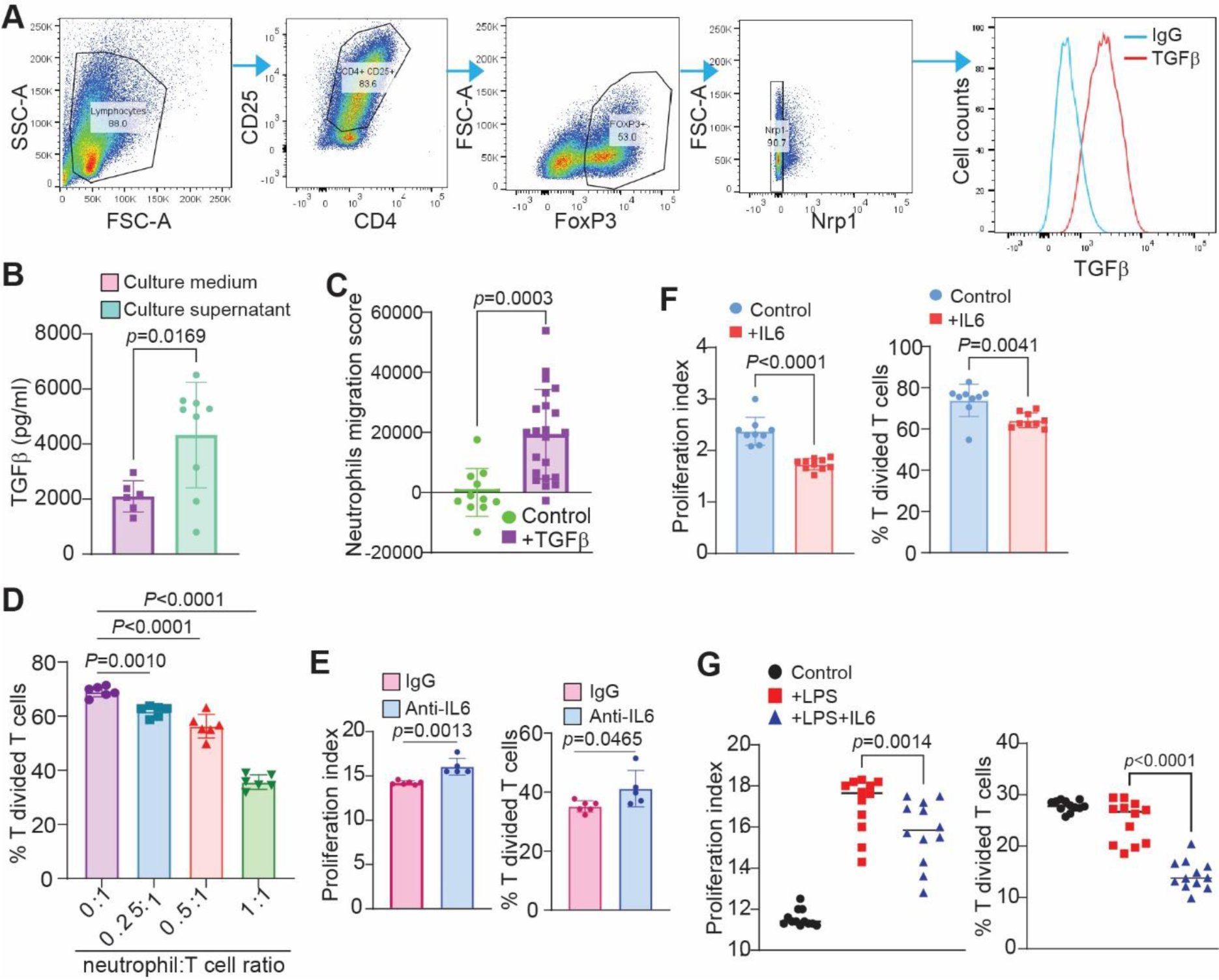
Treg cells produce TGFβ to recruit neutrophils and neutrophils secret IL6 to inhibit activation of T and B cells in mouse colon tumor. **A**. Treg cells express TGFβ. Naïve T cells were isolated from mouse spleens and induced to differentiate into Treg in vitro and analyzed for TGFβ expression by flow cytometry. **B**. Treg cells secret TGFβ in vitro. **C**. TGFβ induces migration of neutrophils in vitro. **D**. Neutrophils inhibit T cell activation in vitro. Neutrophils and T cells were co-cultured at the indicated ratio in anti-CD3/CD28-coated plates for 3 days. **E**. Neutrophils inhibits T cell activation through IL6. Neutrophils and T cells were co-cultured in a 1:1 ratio in the presence of IgG and IL6 neutralization mAb. **F**. IL6 inhibits T cell activation. **G**. IL6 inhibits B cell activation. B cells were cultured in the presence of LPS, ILP and IL6 as indicated and analyzed for B cell proliferation in vitro.

### IL6 blockade immunotherapy suppresses colon tumor growth in vivo

To determine whether the above in vitro findings can be translated to in vivo tumor growth regulation in vivo, we injected CT26 tumor cells to the cecal wall of syngeneic mice to establish orthotopic colon tumor. CT26 is a MSS subtype of colon tumor cell line^69^. The AOM-DSS-induced colorectal tumor is also a DNA mismatch repair proficient MSS colorectal tumor subtype^70^. To block neutrophil function, the CT26 tumor-bearing mice were treated with Ly6G neutralization mAb. Analysis of tumor tissues from the tumor-bearing mice shows that Ly6G blockade therapy effectively depleted neutrophils (Fig. 7A&B). However, neutrophil neutralization did not significantly change CD8^+^ T cell tumor infiltration level (Fig. 7C) and tumor growth (Fig. 7D). We then tested IL6 blockade. Treatment of the CT26 tumor-bearing mice with IL6 neutralization mAb significantly increased tumor-infiltrating CD8^+^ T cells (Fig. 7E) and significantly suppressed tumor growth (Fig. 7F). We therefore conclude that IL6 promotes colorectal tumor immune evasion.

**Figure 7.**
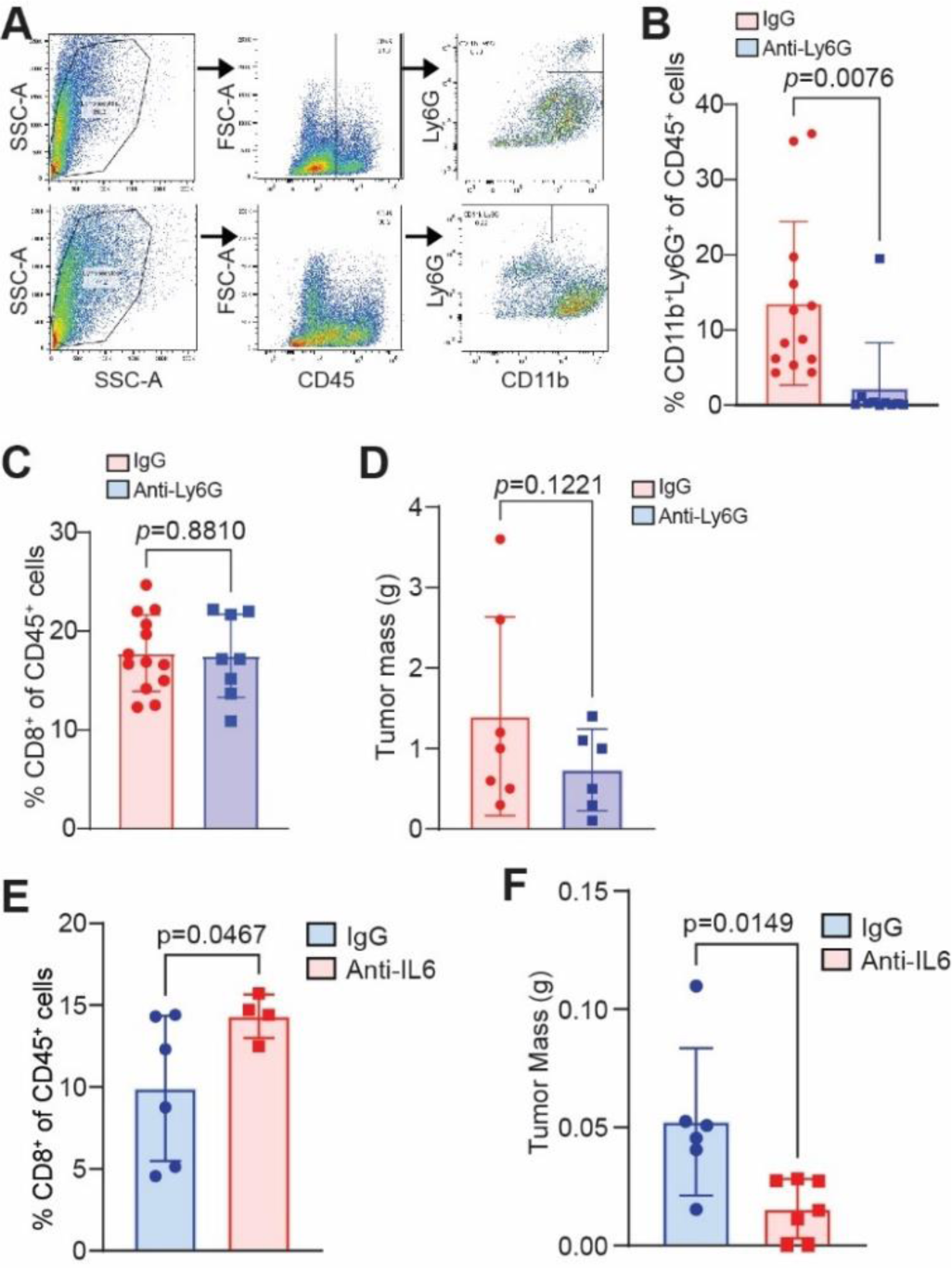
IL6 blockade suppresses colon tumor growth in vivo. **A**. CT26 orthotopic tumor-bearing mice were treated with Ly6G neutralization mAb. Shown is flow cytometry analysis gating strategy of the tumor. **B**. Quantification of neutrophils in tumor as shown in A. **C**. Quantification of CD8^+^ tumor-infiltrating T cells. **D**. Tumor weight. **E**. CT26 orthotopic tumor-bearing mice were treated with IL6 neutralization mAb. Shown is quantification of tumor-infiltrating CD8^+^ T cells by flow cytometry. F. Tumor weight.

### CD40 agonist antibody immunotherapy activates B cells to suppress colon tumor immune evasion in vivo

Global *Cd274* deletion in mice leads to a dramatic decrease in B cells in colorectal tumor in the inflammation-driven colorectal tumor mouse model (Fig. 3C & E). B cells have recently emerged as key regulators of T cell activation in anti-tumor immune response^71–74^. Unique subsets of B cells have been shown to activate T cell activation to suppress tumor growth^74^. We therefore further analyzed B cells in colorectal tumor. UMAP projection analysis indicates that the small population of cells are plasma cells and majority of cells and B cells. Among the B cells, eight cell clusters were identified with an increased subpopulation of Satb1^+^ and decreased subpopulation of CD80^+^ cells in the WT colon tumor (Fig. 8A-C). To functionally determine whether decreased B cells are responsible for the increased colorectal tumorigenesis, we performed a proof-of-concept study. The CD40-CD40L pathway activates B cells into antigen-presenting cells^75^. CD40 agonist mAb is well-documented to activate B cells to prime T cells in an anti-tumor immune response^76–78^. We then used CD40 agonist mAb to treat the CT26 colon tumor-bearing mice. The CD40 agonist mAb-treated mice exhibit splenomegaly with more than three times enlarged spleens (Fig. 8D). However, the percentage of CD45^+^ cells decreased in the enlarged spleens (Fig. 8E & F). Analysis of immune cells in the spleens of tumor-bearing mice indicated that CD40 agonist therapy increased CD4^+^ and CD8^+^ T cells (Fig. 8F). The % B cell level is not significantly changed (Fig. 8F). Because the three-times enlarged spleen size, the absolute B cell number in the spleen increased significantly (Fig. 8F). Analysis of the colon tumor revealed that CD45^+^ cells increased significantly in CD40 agonist mAb-treated mice (Fig. 8G&H). Although the CD4^+^ and CD8^+^ T cells did not increase based on CD45 level, the levels of total tumor-infiltrating CD4^+^ and CD8^+^ T cells are significantly higher in the CD40 agonist mAb-treated mice as compared to the control mice (Fig. 8H). Consistent with increased T cells, CD40 agonist mAb therapy significantly inhibited the growth of the CT26 tumor in mice (Fig. 8I). Taken together, our findings determine that B cells plays a key role in T cell activation and colon tumor growth control and PD-L1 protects B cells in colorectal tumor to suppress colorectal tumorigenesis in vivo.

**Figure 8.**
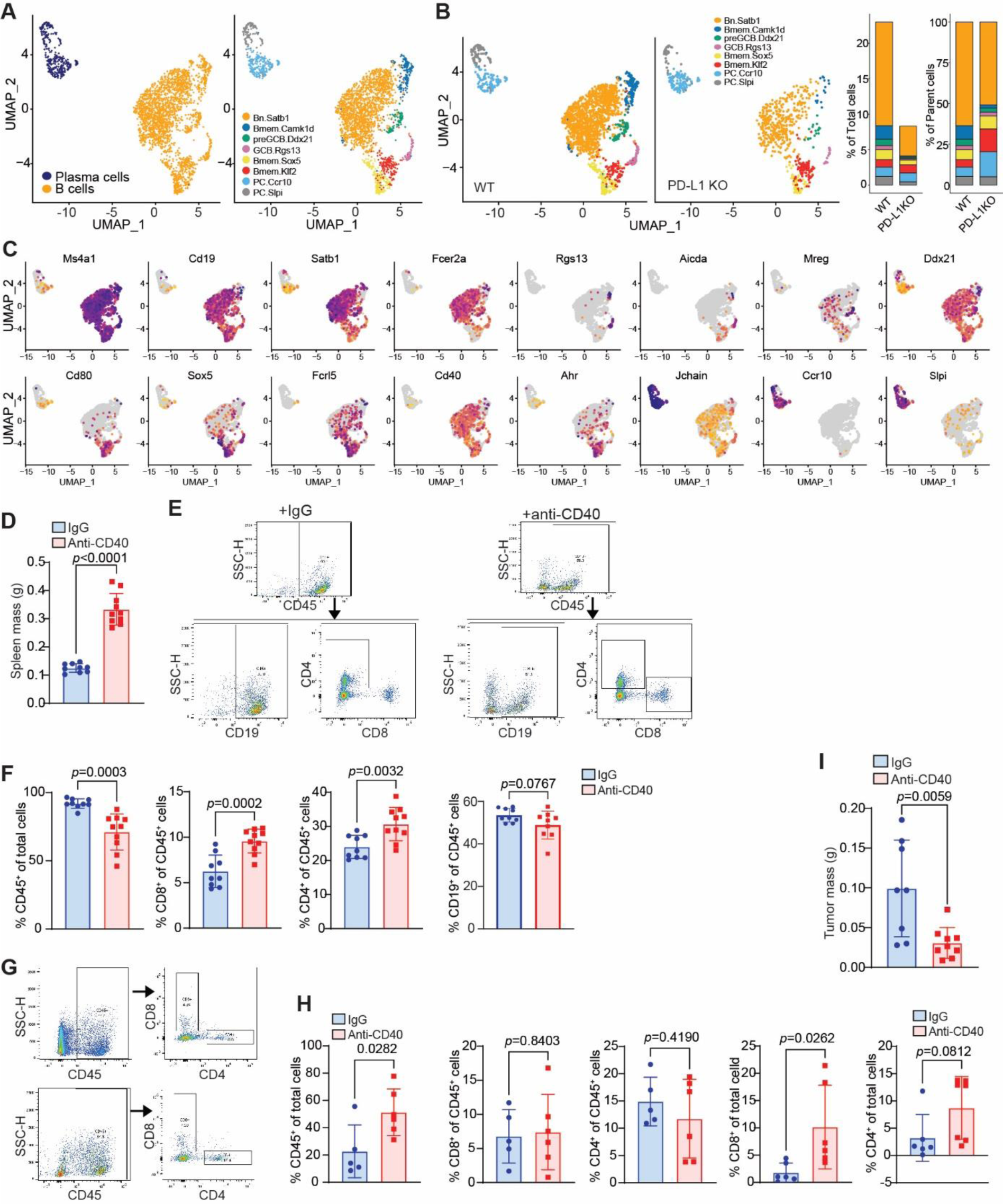
PD-L1 restrains B cell activation and expansion to promote colon tumor immune escape. **A**. UMAP plots of B and plasma cells (left), B cell subpopulations (middle), and B cell subpopulation by genotypes (right). **B**. Barplot of B cell subpopulation as shown in A. **C**. UMAP plots showing the indicated B cell marker gene transcripts in mouse colon tumor. **D**. Spleen weight in CD40 agonist mAb-treated CT26 orthotopic tumor-bearing mice. **E**. Flow cytometry analysis of immune cells of spleens in CD40 agonist mAb-treated CT26 orthotopic tumor-bearing mice. Shown is gating strategy. **F**. Quantification of the indicated cell subsets in spleens of the CT26 orthotopic tumor as analyzed by flow cytometry. **G**. Flow cytometry analysis of immune cells in the CT26 orthotopic tumor. Shown is gating strategy. **H**. Quantification of the indicated cell types in the CD40 agonist mAb-treated CT26 orthotopic tumor. I. Tumor weight.

## Discussion

Colorectal cancer was the first human neoplasia found to be under immunosurveillance more than a decade ago^79^. The type, density and location of tumor-infiltrating immune cells correlate with tumor initiation, progression and recurrence^80–85^. Human colorectal tumors are therefore a type of highly immunogenic cancer^79^. However, human colorectal cancer is the third most commonly diagnosed human cancer, suggesting that the host immunosurveillance failed to detect and eliminate colorectal tumorigenesis despite its high immunogenicity. Furthermore, approximately 85-90% human colorectal cancer is the MSS subtype^17^, whereas MSI-H subtype accounts for only 10-15% of all human colorectal cancer cases. This is an unexpected phenomenon since both tumor cells and tumor-infiltrating immune cells of MSI-H human colorectal cancer express high level of PD-L1, whereas both tumor cells and tumor-infiltrating immune cells of MSS human colorectal cancer express weak to undetectable PD-L1^19,20^. PD-L1 is an immune suppressive ligand that promote tumor immune evasion^1,2^, it therefore would be expected that the PD-L1^hi^ MSI-H colorectal cancer should the dominant human colorectal cancer, but this is also not the case, suggesting that PD-L1 might function as a tumor suppressor in certain colorectal cancer. In this study, we determined that loss of host PD-L1 leads to increased colorectal tumorigenesis in the inflammation-driven MSS colorectal tumor mouse model. Our findings determine that PD-L1, in the context of certain colorectal cancer, functions as a tumor suppressor.

PD-L1 also functions as a suppressor of colonic inflammation^24,25^, a process that promotes colorectal tumor development^14^. It has been proposed that the human colon is in a constant low-grade inflammation state^13,15^, PD-L1 thus simultaneously execute two conflicting functions in the context of colorectal tumorigenesis. Our findings indicate that the function of PD-L1 as a colonic inflammation suppressor and thereby a colorectal cancer suppressor overpowers its function in promoting colorectal tumor immune evasion. This notion is supported by our finding that levels of both CD4^+^ and CD8^+^ T cells are not increased in the tumor in PD-L1 KO tumor-bearing mice as compared to WT mice, suggesting that loss of host PD-L1 does not unleash T cells during colorectal tumorigenesis. The levels of IFNγ and TNFα proteins, two effectors of activated T cells, are also not increased in PD-L1 KO tumor-bearing mice. These findings indicates that PD-L1 function in suppression of T cells is impaired in the inflammatory colonic microenvironment, implying thereby that the main function of PD-L1 in the colon is suppression of colonic inflammation and resultant suppression of colorectal tumorigenesis.

Emerging experimental data points to a crucial role of gut microbiota in colonic inflammation and host cancer immunosurveillance^16,44,52,86^. Gut microbiome metabolites regulate activation and function of both innate and adaptive immune responses to mediate colonic inflammation and tumor immune evasion^16,44,86^. The tryptophan metabolites kynurenine and indoles are key immune regulators that link microbiome to host immune response regulation. Bacteria metabolize tryptophan to indoles and kynurenine^46–48^. Indoles and kynurenine activate AhR to regulate expression of responsive genes in both innate and adaptive immune cells^44,49^. *Lactobacillus* metabolizes tryptophan to generate indoles and likely kynurenine to activate AhR in tumor-bearing hosts^44^. In this study, we observed that abundance of *L. murinus* in the gut is controlled by host PD-L1, and loss of PD-L1 leads to a dramatic increase in gut *L. murinus* in tumor-bearing mice. Our findings determine that PD-L1 is a suppressor of *L. murinus* in the gut and loss of PD-L1 results in increased *L. murinus* and AhR pathway activation to promote colorectal tumorigenesis.

A consequence of AhR pathway activation is enhanced homing and function of Treg cells^53,54,87^. Loss of host PD-L1 therefore may not only activates the *Lactobacillus*-AhR pathway to recruit Treg cells to the colon and tumor^53,54,87^, but may also directly unleash PD-1^+^ Treg cells in the tumor^88–90^. In this study, we observed that loss of PD-L1 leads to increased Treg in colorectal tumor. Treg cells are key suppressor of colonic inflammation^38,91,92^. However, Nrp1 is required for Treg immune suppressive function and stability, and Nrp1^hi^ Treg exhibits higher suppressive activity than Nrp1^lo^ Treg cells^93–95^. In this study, we observed that loss of host PD-L1 increased PD-1^hi^Nrp1^lo^TGFβ^+^Treg cells. The Nrp1^lo^Treg cells may have no significant activity in suppression of colonic inflammation^93–95^. In this study, we determined that Treg cells function as a recruiter and use TGFβ to recruit IL6^+^ neutrophils to the colon tumor microenvironment.

Neutrophils have both pro- and anti-tumor activity^59,60,96^. Consistent with this phenomenon, neutrophil blockade with the Ly6G-specific mAb has no effect on colonic inflammation and colitis^97^. In this study, we determined that neutrophil blockade does not alter colon tumor growth. However, IL6 blockade significantly suppressed colon tumor growth. We further determined that IL6 inhibits B and T cell activation. It is possible that Ly6G^-^ neutrophils may contribute to IL6 production. Further studies are therefore needed to further characterize the neutrophil subpopulations. Nevertheless, our findings determine that host PD-L1 restrains the *Lactobacillus*/AhR/PD-1^hi^Nrp1^li^TGFβ^+^ Treg cells to suppress accumulation of IL6^+^ neutrophils to maintain B and T cell activation and host cancer immunosurveillance to suppress colorectal cancer.

### Limitations of the study

We focused on mechanisms underlying PD-L1 function as a colorectal tumor suppressor in the commonly used the AOM-DSS sporadic and inflammation-driven colorectal tumor mouse model. Given the concept that colon is in a constant low-grade inflammation status, other spontaneous colorectal tumor models are needed to determine whether PD-L1 functions as a suppressor only in inflammation-driven colorectal tumorigenesis, or a general suppressor of the PD-L1^lo^ MSS colorectal tumorigenesis. Furthermore, the mechanism underlying PD-L1 control of *L. murinus* abundance also requires further studies. Nevertheless, our study determines that PD-L1 functions as a tumor suppressor in inflammation-driven colorectal tumorigenesis at least in part via restricting *L. murinus* and the AhR pathway to suppress PD-1^+^Nrp1^lo^TGFβ^+^Treg cells to block IL6^+^ neutrophil tumor recruitment to sustain T and B cell activation and host cancer immunosurveillance.

## Methods

### Mice

*Cd274*^fx/fx^ mice were created as described previously^34^. LyzCre (B6.129P2-Lyz2^tm1^(cre)^Ifo^/J) and the Mrp8Cre (B6.Cg-Tg(S100A8-cre,-EGFP)1Ilw/J) were obtained from the Jackson Laboratory (Bar Harbor, ME). C57BL/6 and BALB/c mice were obtained from the Jackson Laboratory and Charles River Laboratories. Use of mice is approved by the Augusta University institute animal care and use committee.

### The AOM-DSS spontaneous colorectal tumor model

Mice were injected with azoxymethane (AOM) 10 mg/kg body weight, intraperitoneally, once. The day after AOM injection, mice were given dextran sodium sulfate (DSS), 2%, in the drinking water for 1 week. The DSS was then replaced with drinking water for 2 weeks. The DSS was repeated twice for a total of 3 cycles. Mice were maintained on normal drinking water until being sacrificed. Mice were weighed every 3-4 days and checked daily for survival. Colon tissues were harvested and cleaned thoroughly with PBS. The colons were cut open longitudinally. Both male and female mice were used for all genotypes. All mice were genotyped prior to the experiment.

### AOM-induced sporadic colorectal tumor model

Mice were injected with AOM (10 mg/kg body weight) intraperitoneally every week for 5 weeks. They were maintained in standard housing for 143 days before being analyzed for tumor development.

### Tumor cell lines

CT26 cell line was obtained from ATCC (Manassas, VA). MC38-met cells were as previously described ^5^. All cells were cultured in RPMI1640 medium with 10% FBS at 37^0^C in 5% CO2 incubator.

### Tumor cell transplant

Tumor cells were cultured in RPMI160 with 10% FBS for 24 h, and then harvested with trypsin and washed three times with PBS to remove residual media. Tumor cells were resuspended in PBS and injected subcutaneously into the right flank, or orthotopically into the cecal wall of mice.

### In vivo blockade

Tumor cells were implanted orthotopically to the cecum using an insulin syringe (5×10^4^ cells/mouse). Treatment began on days 3-10. Mice received 200 μg of the blockade mAbs every 3 days via ip injection.

### scRNA-sequencing

Colons were collected from the tumor-bearing mice. Tumor nodules were dissected from the colon and digested with collagenase/hyaluronidase/DNaseI solution to RPMI1640 medium at 37°C for 30 minutes with continuous agitation by stir bar and needle aspiration. Live cells were subsequently isolated by lymphocyte separation medium gradient centrifugation. Single cell isolation and library generation were performed by 10xGenomic protocol. Analysis was conducted with Seurat (v3) in R. Cells were subsetted to those with <15% mitochondrial reads and 200-6000 RNA features and integrated with Harmony (v0.1). Doublets were identified and discarded using DoubletFinder. Cells then underwent nearest-neighbor clustering; clusters were annotated by CelliD and manual review. Single-cell RNA sequencing reads were mapped to the mouse reference genome and processed into gene expression matrices with CellRanger (l0x Genomics; version 2.1.1, 2.2.0, 3.0.2 and 3.1.0).

### Tumor and colon digestion

Tumors were manually dissected and digested using a collagenase/hyaluronidase/DNase I solution in scintillation vials with magnetic stir rods to break up the tissue. Tissues are digested for 25 minutes at 37°C or at room temperature for 1 hour.

### Flow cytometry

Cells resuspended in PBS or FACS buffer. Antibodies were added to appropriate concentration and stained at 4°C for 30-60 min, then washed with PBS+0.5% BSA. Cells were fixed in in 2% paraformaldehyde. For Zombie UV staining: Cells resuspended in PBS or FACS buffer. Zombie UV (1:1000) added, incubate for 10 minutes at room temperature in the dark. Antibodies then added at appropriate concentrations and stain as above. Intracellular staining with BD Cytofix/Cytoperm kit (catalog 555028). Cells were resuspended in FACS buffer with Golgiplug for 2 hours. Stained for surface markers per above protocol. Washed 2x with FACS buffer. Resuspended in permeabilization solution, then washed 2x in perm/wash buffer. Resuspended in perm/wash buffer. Antibodies for intracellular proteins of interest added, incubate for 20 minutes at room temperature in the dark. Washed in perm/wash buffer, resuspended in 2% paraformaldehyde.

### Immunohistochemistry

The slides were dewaxed using xylene and rehydrated in 100%, 90%, 70%, and 50% ethanol sequentially. Antigen retrieval was performed using antigen unmasking solution and antigen retrienval solution, pH 6 or pH 9. Slides were then blocked, and the antibody was added. Next, the slides were washed 3 times in TBST. Next, the Opal flurophores were added for 10 minutes. The slides were again washed, and then rinsed in antigen retrieval solution (pH 6 or pH 9). This process was repeated for each antibody, sequentially. Lastly, the slides were stained with DAPI and mounted with slide covers.

### Analysis of colonic inflammation

Colon tissues were prepared as Swiss rolls and fixed in 10% formalin overnight. The fixed tissues were processed into paraffin blocks and cut into 10 micron sections. For the inflammation score, each grade represents the following: Grade 0: normal colonic mucosa; Grade 1: loss of 1/3 of the crypts; Grade 2: loss of 2/3 of the crypts; Grade 3: lamina propria is covered with a single layer of epithelium, mild inflammatory cell infiltrate present; Grade 4: erosions and marked inflammatory cell infiltration present. The slides were evaluated by two board certified pathologists.

### ELISA

A 96-well assay plate was coated overnight with capture antibody. The plate was then washed 3 times with 0.05% Tween-20 in PBS and blocked for 1-3 hours with 1x assay diluent. Standards and samples diluted as appropriate in assay diluent were then added to the wells, in duplicate/triplicate. The plate was then incubated 1-3 hours, and washed 3 times with wash buffer. Capture antibody was then added for 1 hour, and the plate was again washed times. Avidin-HRP was added for 30 minutes, and then washed 3 times. 1:1 TMB substrate was then added to each well, and the plate was left to incubate in the dark for 10-15 minutes (based on manufacturer protocol). The reaction was stopped with 1M H_2_SO_4_ and plate was read for absorbance in a plate reader.

### Serum cytokine analysis

Blood samples were collected either via cardia bleed or via a submandibular vein bleed into serum gel tubes. Tubes were allowed to clot for at least 30 minutes, then centrifuged to collect the serum. Simultaneous quantification of cytokines in murine serum was performed using LEGENDplex Mouse Inflammation Panel (TNF-α, IFN-γ, IL-1a, IL-1b, IL-6, IL-10, IL-17A, IL-12p70, GM-CSF, IL-23, IFN-b, MCP-1, IL-27; BioLegend Cat# 740446), Mouse TH Panel v4 (TNF-α, IFN-γ, IL-4, IL-2, IL-6, IL-10, IL-17A, IL-17F, IL-5, IL-9, IL-13, IL-22; BioLegend Cat# 741044) and Mouse Custom696 Panel (CXCL1(KC), CCL3(MIP-1a), CCL5(RANTES), CCL20(MIP-3a), CCL11(Eotaxin), TGFb1(FreeActive), CXCL10(IP-10), CXCL9(MIG), VEGF, CCL4(MIP-1b), IFN-a CXCL12(SDF-1); BioLegend Cat# 900001796) according to manufacturer’s instructions. In brief, serum samples were diluted 2-fold with assay buffer and standards were mixed with matrix solution (Biolegend) to account for additional components in the serum samples. Standards and samples were plated with capture beads and incubated for 2 h at room temperature on plate shaker (800 rpm). After washing the plate with wash buffer, detection antibodies were added to each well. The plate was incubated on shaker for 1h at room temperature. Finally, without washing, SA-PE was added and incubated for 30 min. Samples were acquired on Novocyte Quanteon flow cytometer (Agilent Technologies). Standard curves and protein concentration were calculated using R package DrLumi installed on R 3.5.2. The limit of detection was calculated as an average of background samples plus 3xSD. Assay and data calculations were performed at Immune Monitoring Shared Resource laboratory.

### T cell proliferation assay

CD3^+^ T cells were purified from BALB/c mouse spleen cells with the MojoSort mouse CD3 T cell isolation kit (Biolegend, San Diego, CA) according to the manufacturer’s instructions. Isolated cells are incubated in a 37°C water bath in prewarmed PBS+0.15-0.30μM CFSE (Life Technologies, Carlsbad, CA, USA)for 15 minutes, vortexing every 5 minutes. The cells are then washed and incubated in RPMI 1640 medium with 10% FBS at a volume of 5x the volume used for labelling for 30 minutes, vortexing every 10 minutes. The cells are again washed 2 times with medium and plated. CFSE intensity is analyzed using a flow cytometer. For T cell proliferation assay, a 96-well culture plate was coated with anti-mouse CD3 (8 μg/mL), anti-mouse CD28 MAbs (10 μg/mL), and re. The purified T cells were labeled with and then seeded in the plate at a density of 1.5 × 105 cells/well in 150 μL medium for 3 days. Cells were analyzed by flow cytometry.

### Neutrophil isolation and migration assay

Neutrophils were isolated using the StemCell neutrophil isolation kit or the Biolegend Neutrophil isolation kit, per manufacturer protocols. Neutropihls were isolated from the spleen of a 4T1 tumor bearing mouse. The QCM migration plate was brought to room temperature. 250 μL of cells (2×10^6^/well) were added to each well insert. 400μL of serum-free media and TGFβ was added to the lower chamber. The cells were incubated for 18-24 hours. On day 2, 225 μL of the lower well was transferred to a white walled 96 well plate for fluorescence after following manufacturer instructions for adherent cells. The CyQuant dye was diluted 1:75 with 4x lysis buffer. 75 μL of the dye/lysis buffer mixture was added to each well of the 96 well plate. Flourescence was read on 48/520nm channel. For neutrophil migration with Treg in lower chamber: follow manufacturer protocol to release adherent cells, then harvest all of the lower chamber and transfer to flow cytometry tube. Stain for CD11b and CD3. Quantify changes in CD11b cells.

### Treg differentiation

24-well plates were coated with anti-CD3 mAb (8 μg/ml) overnight in 4°C. Single cell suspension was prepared from mouse spleen and CD4^+^ cells were isolated using Biolegend CD4 isolation kit. Treg were differentiated from the purified CD4^+^ cells using CellXVivo Mouse Treg differentiation kit (R&D System). Briefly, cells were suspended in Treg differentiation media, then added to the anti-CD3 mAb-coated plate. The cells were incubated for 5 days, then analyzed via flow cytometry.

### B cell activation

B cells were isolated from naive BALB/c mouse spleens using the BioLegend Pan-B cell isolation kit. The cells were then labelled with CFSE. Cells were stimulated with lipopolysaccharise (1μg/mL).

### 16S rRNA gene sequencing and shotgun metagenomics sequencing

Fecal samples were collected from the cecum of mice using aseptic technique. Samples were processed and analyzed with the ZymoBIOMICS Targeted Sequencing Service (Zymo Research, Irvine, CA). The ZymoBIOMICS®-96 MagBead DNA Kit (Zymo Research, Irvine, CA) was used to extract DNA using an automated platform. Bacterial 16S rRNA gene targeted sequencing was performed using the Quick-16S NGS Library Prep Kit (Zymo Research, Irvine, CA). The bacterial 16S primers amplified the V3-V4 region of the 16S rRNA gene. The sequencing library was prepared using an innovative library preparation process in which PCR reactions were performed in real-time PCR machines to control cycles and therefore limit PCR chimera formation. The final PCR products were quantified with qPCR fluorescence readings and pooled together based on equal molarity. The final pooled library was cleaned with the Select-a-Size DNA Clean & Concentrator (Zymo Research, Irvine, CA), then quantified with TapeStation (Agilent Technologies, Santa Clara, CA) and Qubit (Thermo Fisher Scientific, Waltham, WA). The ZymoBIOMICS Microbial Community Standard (Zymo Research, Irvine, CA) was used as a positive control for each DNA extraction. The ZymoBIOMICS Microbial Community DNA Standard (Zymo Research, Irvine, CA) was used as a positive control for each targeted library preparation. Negative controls (i.e. blank extraction control, blank library preparation control) were included to assess the level of bioburden carried by the wet-lab process. The final library was sequenced on Illumina MiSeq with a v3 reagent kit. The sequencing was performed with 10% PhiX spike-in. Unique amplicon sequences variants were inferred from raw reads using the DADA2 pipeline. Potential sequencing errors and chimeric sequences were also removed with the Dada2 pipeline. Chimeric sequences were also removed with the DADA2 pipeline. Taxonomy assignment was performed using Uclust from Qiime v.1.9.1 with the Zymo Research Database. Composition visualization, alpha-diversity, and beta-diversity analyses were performed with Qiime v.1.9.1. Taxonomy that have significant abundance among different groups were identified by LEfSe using default settings. Other analyses such as heatmaps, Taxa2ASV Deomposer, and PCoA plots were performed with internal scripts. The number of genome copies per microliter DNA sample (genome copies) was calculated by dividing the gene copy number by an assumed number of gene copies per genome.

### Metabolomics

Samples were extracted in extraction solution (CAN: Methanol=1:4, V/V) containing internal standards and used for LC-MS analysis. The data acquisition instruments consisted of Ultra Performance Liquid Chromatography (UPLC) and Quadrupole-Time of Flight Spectrometry (TripleTOF 6600+, AB SCIEX). The data acquisition instruments consisted of Ultra Performance Liquid Chromatography (UPLC) and tandem mass pectrometry (MS/MS) (QTRAP®6500+). LIT and triple quadrupole (QQQ) scans were acquired on a triple quadrupole-linear ion trap mass spectrometer QTRAP LC-MS/MS System, equipped with an ESI Turbo Ion-Spray interface, operating in positive and negative ion mode and controlled by Analyst 1.6.3 software (Sciex). The ESI source operation parameters were as follows: source temperature 500°C; ion spray voltage (IS) 5500 V (positive), -4500 V (negative); ion source gas I (GSI), gas II (GSII), curtain gas (CUR) were set at 50, 50, and 25.0 psi, respectively; the collision gas (CAD) was high. Instrument tuning and mass calibration were performed with 10 and 100 μmol/L polypropylene glycol solutions in QQQ and LIT modes, respectively. A specific set of MRM transitions were monitored for each period according to the metabolites eluted within this period.

The mixed samples first underwent untargeted metabolomics detection. Metabolites were analyzed qualitatively with in-house database MWDB, integrated public database, AI database, and MetDNA. The identified metabolites were integrated with the in-house database MWDB (Metware Biotech). Lastly, quantification using MRM mode was performed for all samples based on the newly integrated database. Metabolites were quantified by triple quadrupole mass spectrometry with multiple reaction monitoring (MRM). In MRM mode, the first quadrupole screens the precursor ions for the target compound and excludes ions of other molecular weights. After ionization induced by the impact chamber, the precursor ion is fragmented, and a characteristic fragment ion is selected through the third quadrupole and excludes the interference of other untargeted ions. The samples are classified according to their features such that highest homogeneity is achieved between sample from the mouse group and highest heterogeneity is achieved between samples from different groups. The compound quantification data was normalized (Unit Variance Scaling, UV Scaling) and heatmaps were drawn by R software Pheatmap package. Hierarchical Cluster Analysis (HCA) was used to cluster the samples.

### Statistical analysis

Statistical analysis was performed using GraphPad prism software. Statistical comparisons were conducted using unpaired two-tailed Student’s t test with *p* value < 0.05 being considered as statistically significant.

