## Supplemental Data for "PD-L1 Restrains the PD-1^hi^Nrp1^lo^TGFβ^+^ Treg to block IL6^+^ Neutrophil tumor infiltration to Suppress Inflammation-driven Colorectal Cancer"

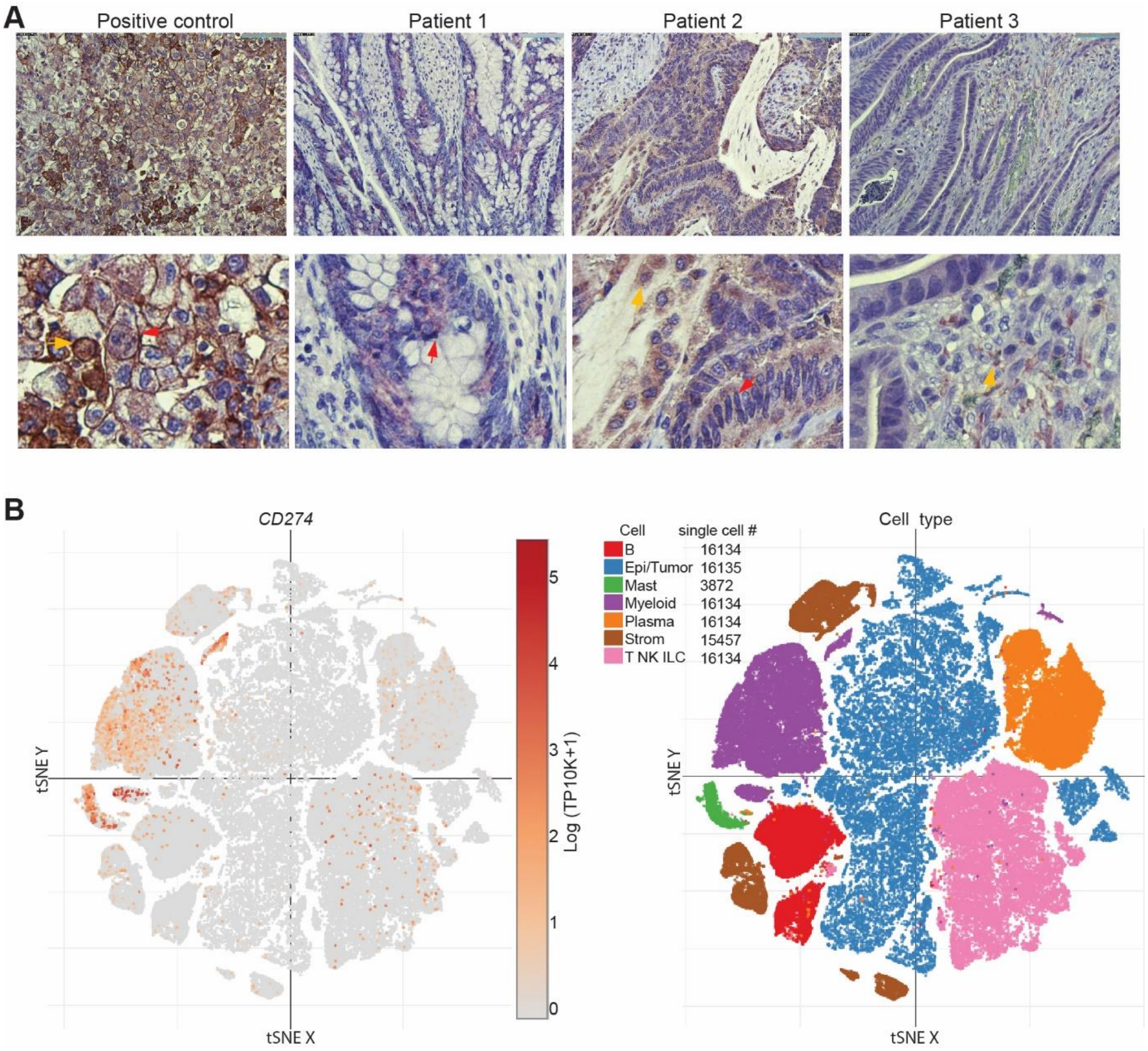


**Figure S1. PD-L1 expression profiles in human colon cancer**. **A**. Immunohistochemical staining of human colon tumor tissues from three patients. Human adrenal tumor tissues were stained as PD-L1 positive control. Shown are low (top) and high (bottom) magnification images. The yellow arrow indicates leukocytes, and the black arrow indicates tumor cells. **B**. PD-L1 protein level in major cell types of human colon cancer in the single-cell level (left) and the corresponding cell types (right). scRNA-seq datasets (GEO:GSE:178341) were analyzed at the Broad Institute's Single Cell Portal.

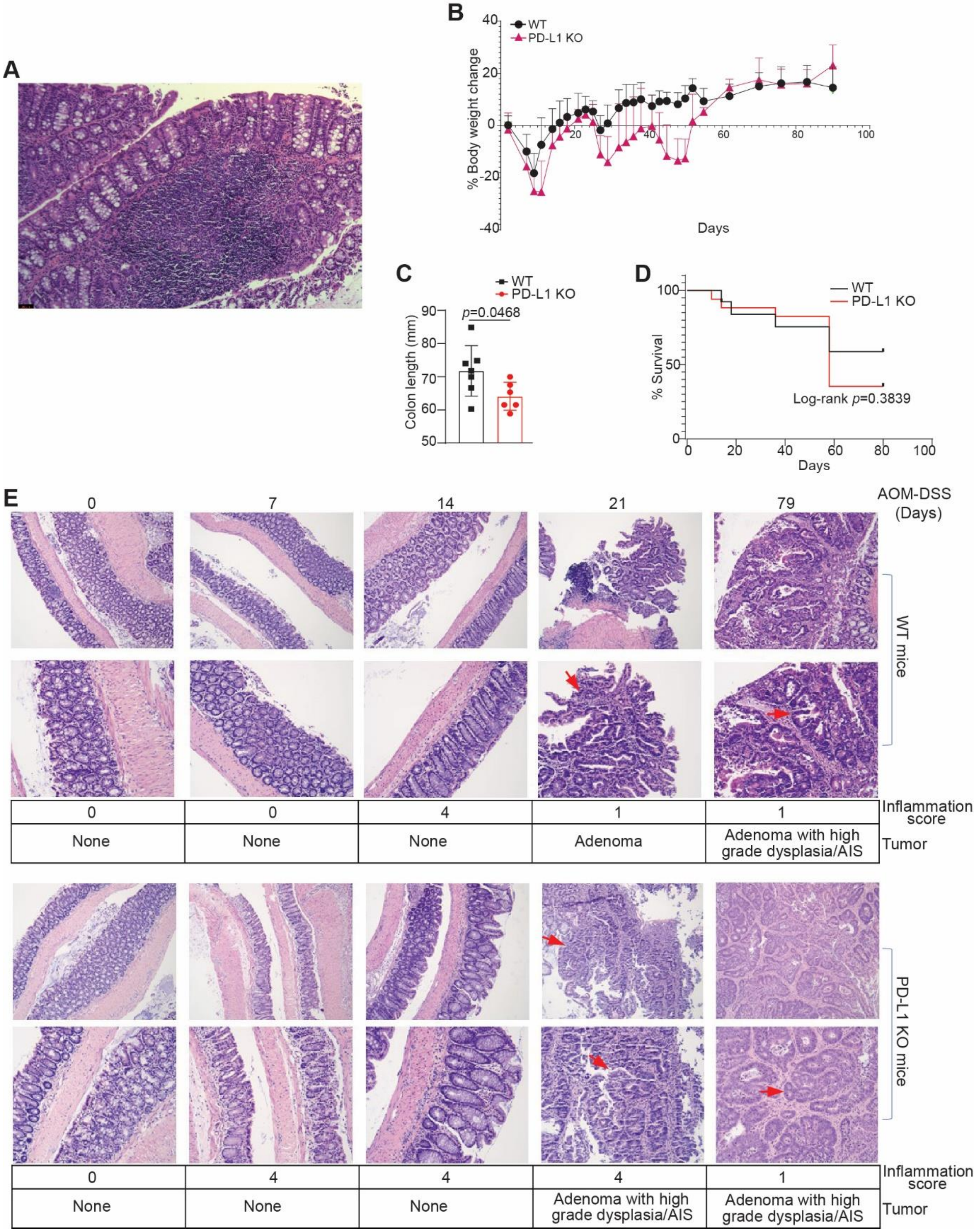


**Figure S2. PD-L1 suppresses colonic inflammation during colon tumorigenesis. A**. A representative colon tissue section from mice treated with five cycles of AOM without DSS. **B**. Mouse body weight change kinetics after AOM-DSS treatment. **C**. Colon length at the end point of AOM-DSS treated mice. **D**. Mouse survival after AOM-DSS treatment. **E**. Colon tissues at the indicated time points were stained by H&E and analyzed for inflammation and tumor development. Bottom panels are magnified images of the top panels in both WT and PD-L1 KO panels.

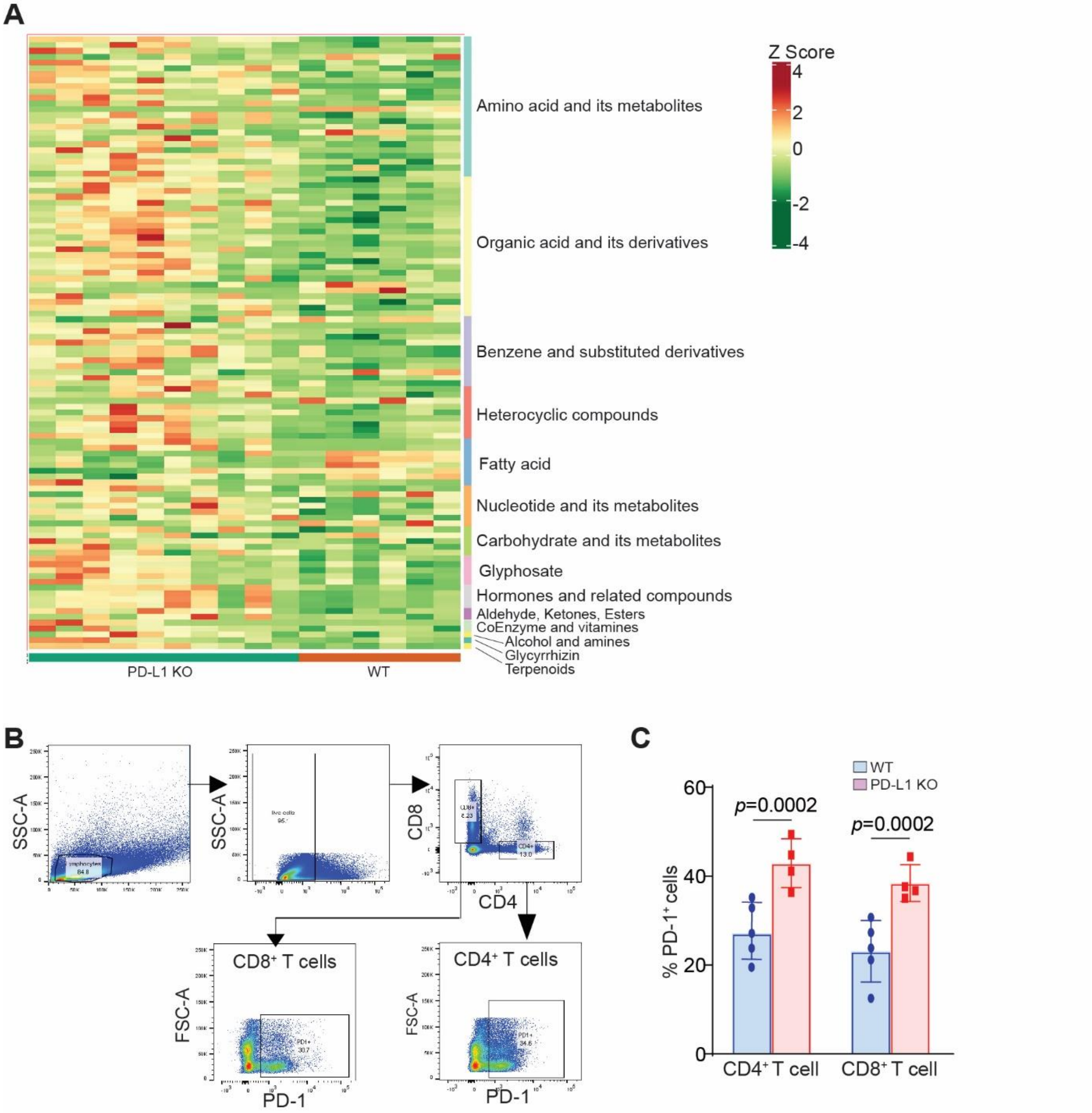


**Figure S3. PD-1 expression in T cells**. **A**. Heatmap of serum metabolites from WT and PD-L1 KO colorectal tumor-bearing mice. Shown is fold-change of differential metabolites quantification using unit variance scaling and plotted the results using ComplexHeatmap in R. **B**. Spleen cells of WT and PD-L1 KO tumor-bearing mice were analyzed by flow cytometry. Shown are gating strategies. **C**. Quantification of PD-1 protein level (MFI) as shown in A.

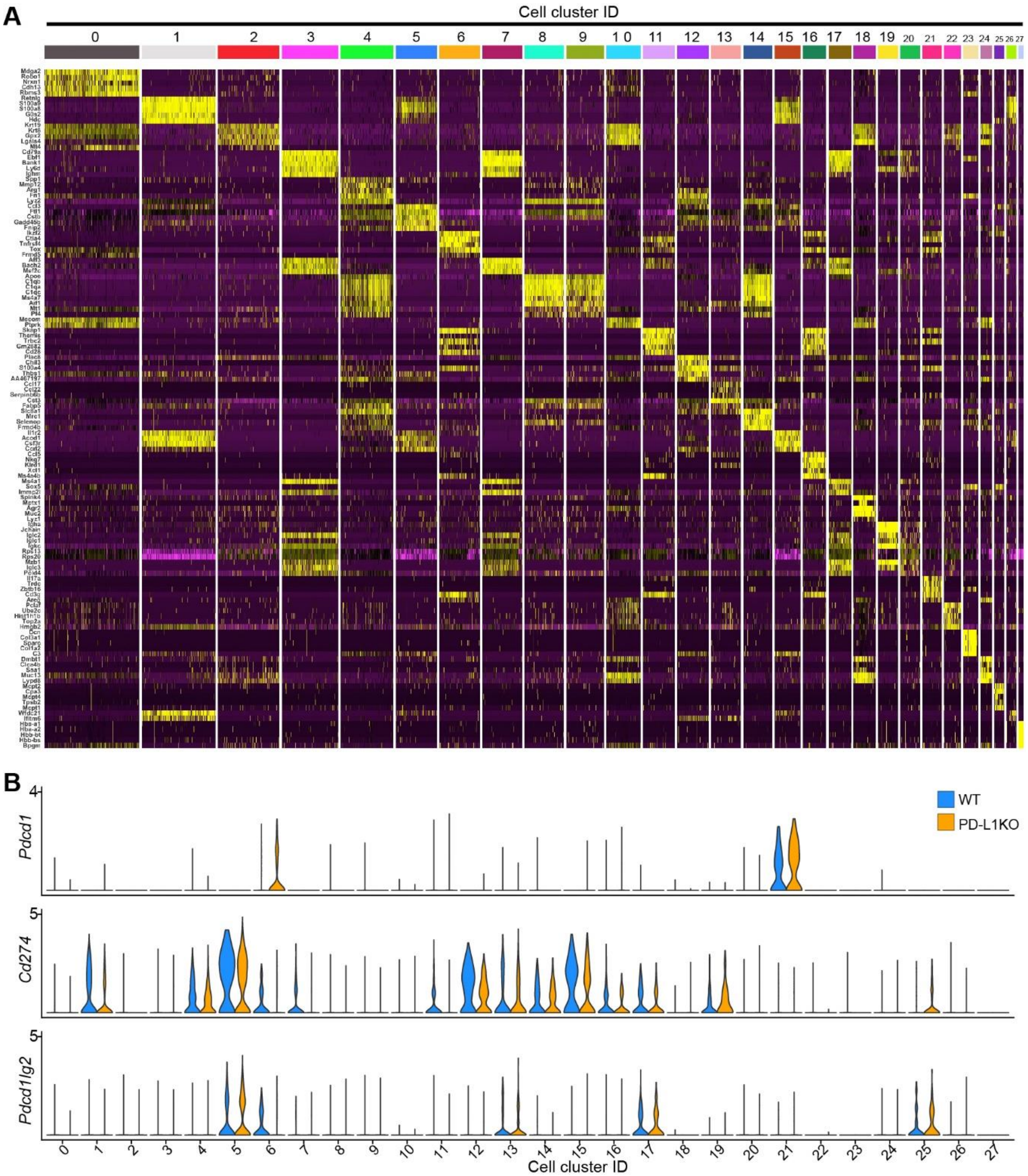


**Figure S4. Cell clustering of AOM-DSS-induced colorectal cancer. A**. Heatmap of the cell cluster signature genes. **B**. Violin plots of the indicated genes in each cell cluster.

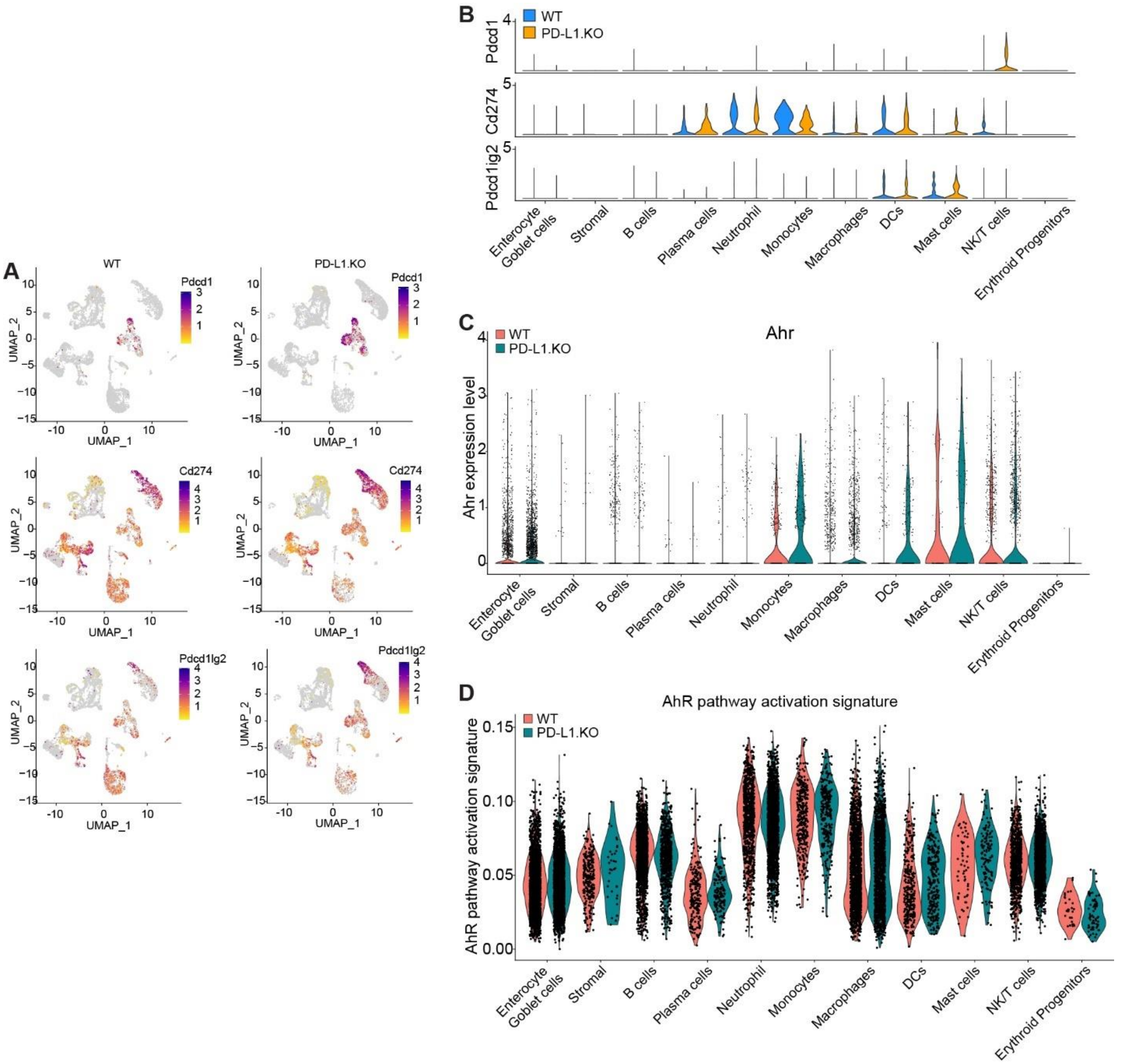


**Figure S5. Immune checkpoint expression in immune cells and AhR pathway activation in colorectal tumor at the single-cell level in WT and PD-L1 KO tumor**. A. UMAP of the indicated genes. **B**. Violin plots of the indicated genes in indicated cell types. **C**. AhR expression level in the indicated immune cell subpopulation in colorectal tumor. **D**. AhR pathway activation signature in the cell subpopulation in colorectal tumor.

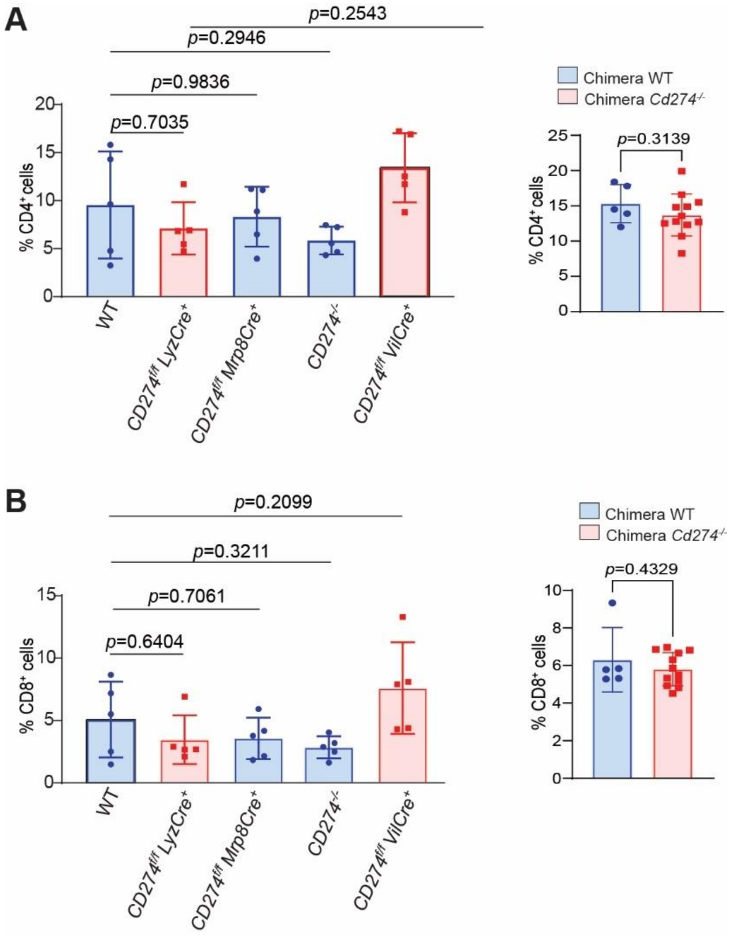


**Figure 6. T cell profiles in PD-L1-deficient tumor-bearing mice**. Mice were treated with AOM-DSS to induce colorectal tumor. Spleen cells were analyzed for $CD4^+$ (A) and $CD8^+$ (B) T cells.

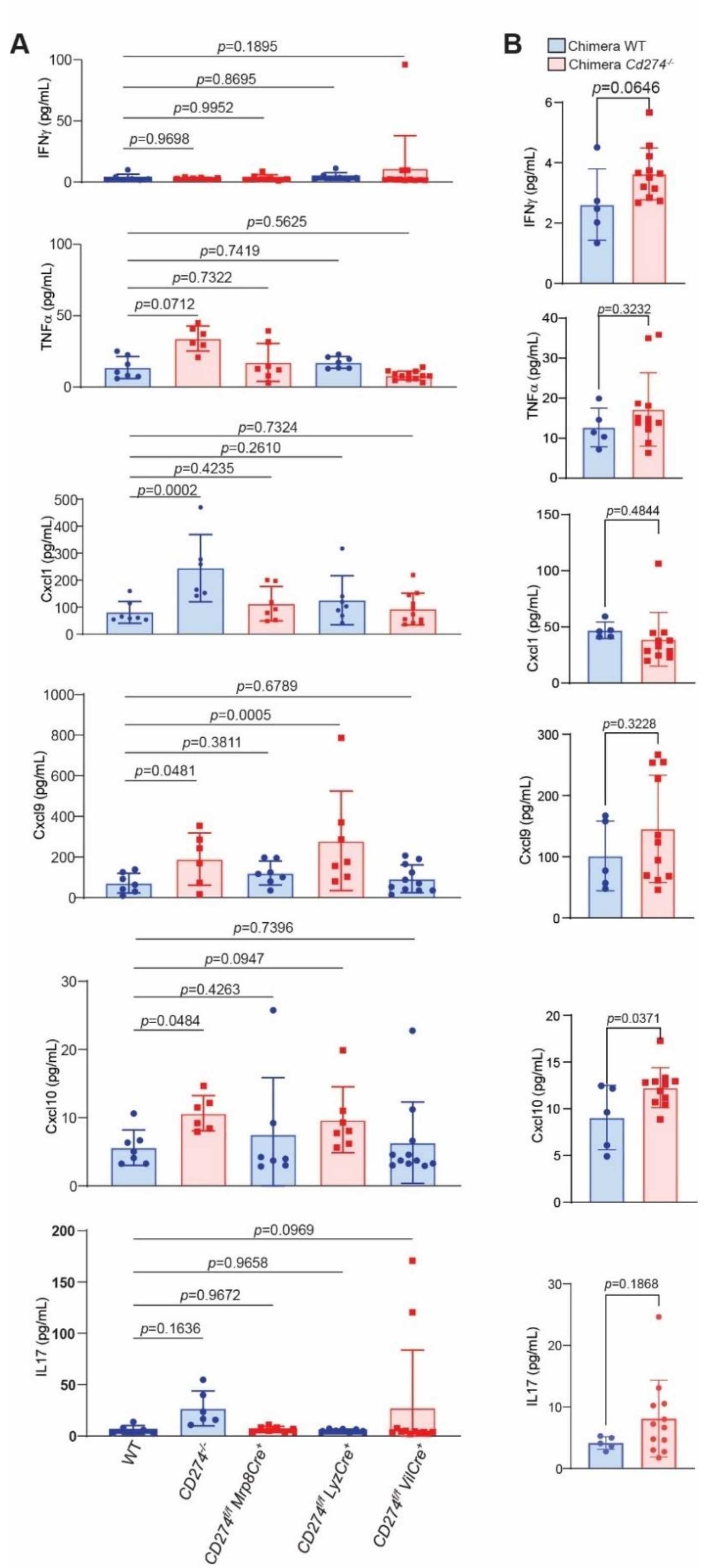


**Figure 7. Cytokine profiles in WT and PD-L1 KO tumor-bearing mice**. Serum was collected from WT and PD-L1 KO tumor-bearing mice and analyzed for the indicated cytokines in WT and the indicated PD-L1 Ko mice (A) and PD-L1 KO chimera mice (B).
